# Inhibition of integrin αvβ8-mediated TGFβ activation and active-TGFβ blockade promote anti-tumor immunity through distinct biological mechanisms

**DOI:** 10.64898/2026.07.06.735099

**Authors:** Katherine Williams, Stephanie Mittman, Natalie S. Firmino, Jasmine W. Larrick, Zhe Zhang, Caroline Whitty, Hsiao-Yen Ma, Xin Ren, Cecilia Chiu, Yagai Yang, Jianhuan Zhang, Minh Thai, Helena Paidassi, Matthieu Masureel, Kelly M. Loyet, Wei-Ching Liang, James T. Koerber, Rafael Cubas, Yan Wu, Shannon J. Turley, Ira Mellman, Nathaniel R. West, Soren Muller, Yan Qu, Dean Sheppard, Alessandra Castiglioni

## Abstract

Transforming Growth Factor β (TGFβ) is a potent immunosuppressor and a primary driver of resistance to cancer immunotherapy. While preclinical models have long suggested that TGFβ inhibition could synergize with immune checkpoint inhibitors, these effects have proven difficult to replicate in clinical settings. The highly regulated TGFβ pathway can be inhibited through various mechanisms, including neutralizing activated ligands or inhibiting upstream activators, such as integrins. Recent structural data demonstrated that integrin αvβ8 can enable TGFβ1/3 signaling without releasing the active cytokines from their Latency-Associated Peptides, suggesting that ligand-blocking antibodies may have limited access to their epitopes. Here, we show that integrin αvβ8 blockade, while achieving anti-tumor responses similar to those of anti-TGFβ antibodies, does so through a distinct mechanism of action. Anti-αvβ8 is 3 orders of magnitude more potent at inhibiting αvβ8-mediated TGFβ activity than a commonly used antibody against the mature form of the cytokine. Whereas TGFβ ligand inhibition has little effect on TGFβ signaling in tumor-draining lymph nodes (tdLN) and requires IFNγ for its anti-tumor effects, αvβ8 blockade strongly inhibits TGFβ signaling in tdLN and, in combination with PD-L1 blockade, drives tumor control through an IFNγ -independent mechanism that strictly requires T cell egress from tdLN. Combined αvβ8 and anti-PD-L1 blockade enhances antigen presentation in dendritic cells (DCs) and, unlike TGFβ ligand blockade, improves the efficiency of DC-induced T cell activation in response to cross-presented antigen. These findings suggest that αvβ8 blockade can disable an immunologically critical source of TGFβ signaling that is not addressed by antibodies targeting TGFβ ligands, suggesting a promising new approach to TGFβ pathway modulation.

**One Sentence Summary:** Unlike ligand-neutralizing antibodies, αvβ8 blockade suppresses TGFβ in tdLN and boosts DC-T cell activation, a differentiated immunotherapy strategy.

## Introduction

Transforming Growth Factor β (TGFβ) inhibition has been shown to amplify cancer immunotherapy (CIT) response in preclinical settings^1–8^. Despite the substantial amount of preclinical data demonstrating synergistic anti-tumor responses when TGFβ pathway inhibition is combined with CIT^1–8^, current attempts to replicate these effects in the clinic have not been satisfactory.

The TGFβ pathway is highly regulated and context-dependent^9^. The three isoforms (TGFβ1, TGFβ2, and TGFβ3) are produced by several cell types in inactive precursor forms in which the active cytokines are associated with their specific Latency-Associated Peptides (LAPs), which prevent binding to the TGFβ receptor. All three can be activated by a diverse set of processes, including proteolytic digestion of the latent complex, interaction with integrins (αvβ6, αvβ8, and αvβ1), changes in pH, interaction with thrombospondin 1, or radiation^10–13^. Once activated, all three TGFβ isoforms can bind to TGFβ receptors to induce phosphorylation of the transcription factors SMAD2 and SMAD3, which then interact with cytosolic SMAD4 to facilitate translocation to the nucleus and regulation of expression of downstream genes^9^.

Most TGFβ-targeting strategies in clinical trials aim to neutralize TGFβ ligands by using antibodies that bind the active cytokines, various forms of ligand traps (TGFΒR-Fc that binds to the mature, active, form of the cytokine), or small molecules that inhibit the kinase activity of the TGFβ receptor (ALK5 inhibitors [ALK5i]). Most of these molecules have a narrow therapeutic window, significant safety concerns, and/or modest anti-tumor efficacy^14–20^. We therefore decided to explore alternative routes to inhibit the TGFβ pathway, aiming to achieve a broader therapeutic window and improved anti-tumor response.

αvβ8 integrin is known to play an important role in activating TGFβ1 and TGFβ3^21,22^. By binding to the TGFβ latent complex presented on cell surfaces (by anchor proteins like GARP/LRRC32 or LRRC33) or bound to the extracellular matrix (by Latent TGFβ Binding Proteins), αvβ8 integrin can activate TGFβ1 and TGFβ3 by altering the conformation of the tethered latent complex, to allow the active TGFβ to be released from the corresponding LAP and to bind to TGFBR2 receptors on adjacent cells. More recently, a new modality has been described by which the αvβ8 integrin activates TGFβ by allosterically changing the structure of the latent complex without releasing the active cytokine^23^.

Previous studies have demonstrated that both anti-TGFβ^4,24^ and anti-αvβ8 blocking antibodies^25^ can augment the effects of checkpoint inhibitors in multiple syngeneic solid tumor models that are generally resistant to CIT. These results have been interpreted as suggesting that the biological effects of anti-TGFβ and anti-αvβ8 antibodies are largely equivalent. However, the recent structural observation that αvβ8-mediated TGFβ activation does not require the release of free TGFβ from the latent complex raises the possibility that TGFβ-blocking antibodies that bind the free, active form of TGFβ might have limited ability to inhibit this alternative signaling mechanism due to steric constraints, and, therefore, anti-αvβ8 integrin and anti-active TGFβ antibodies may have distinct biological effects.

Furthermore, the αvβ8 integrin expression pattern is particularly appealing from a therapeutic perspective. αvβ8 is expressed not only by epithelial cells but also by immune cells that play fundamental roles in regulating immunological responses, including T regulatory cells (T_REG_s) ^26–28^, dendritic cells (DCs) ^29–32^, and monocytes ^33^. Compared with the three TGFβ ligands, which are broadly expressed in a wide variety of healthy and diseased tissues and play multiple roles in tissue homeostasis^34^, αvβ8 expression is restricted to a smaller subset of cells, possibly limiting systemic toxicities observed with more global TGFβ inhibition.

In this study, we set out to compare in more detail the effects of αvβ8-blocking antibodies and a commonly used antibody that blocks all 3 active TGFβ isoforms to test the hypothesis that these two therapeutic strategies might modulate responses to checkpoint inhibitors through distinct biological effects. Our *in vitro* data confirm that the TGFβ ligand-blocking antibody potently inhibits the effects of free TGFβ, whereas the αvβ8-blocking antibody does not. In contrast, the αvβ8 blocking antibody is 3 orders of magnitude more potent in inhibiting αvβ8-mediated TGFβ-activation than the TGFβ blocking antibody. *In vivo*, the αvβ8-blocking antibody is highly efficient at inhibiting the TGFβ pathway in both the tumor-draining lymph nodes (tdLNs) and the tumor microenvironment (TME), as measured by SMAD phosphorylation. In contrast, the TGFβ-blocking antibody inhibits SMAD phosphorylation only in the tumor. Additionally, the efficacy of the TGFβ blocking antibody is completely inhibited by neutralizing IFNγ, and is partially inhibited by inhibition of T cell egress from lymph nodes (by FTY720), whereas the αvβ8 blocking antibody does not require IFNγ, but does require lymph node egress. Single cell RNA sequencing (scRNA-seq) demonstrates that the αvβ8 blocking antibody induces unique molecular phenotypes in tumor conventional DCs (cDCs), upregulating expression of antigen-presentation-related genes, and significantly reducing the presence of CCR7+ migratory DCs. Finally, tdLN DCs sorted from mice treated with an αvβ8-blocking antibody, together with anti-PD-L1, demonstrate enhanced antigen presentation by CD8 T cells, a phenomenon not observed in DCs from mice treated with the TGFβ-blocking antibody plus anti-PD-L1. Together, these results suggest that αvβ8 inhibition enhances the effects of checkpoint blockade through mechanisms that are distinct from the ones mediated by TGFβ ligand-neutralizing antibody.

## Results

### *In vitro*, active TGFβ-neutralizing antibody shows limited activity in blocking αvβ8-dependent TGFβ activation

Given recent observations suggesting that integrin αvβ8 could activate the TGFβ pathway without releasing active TGFβ from the latent complex^23^, we wondered whether an anti-TGFβ antibody^5,24^ known to bind all three active TGFβ isoforms would effectively block TGFβ from binding to TGFBR2 in this context. To gain a deeper structural understanding of the assembly and accessibility of the anti-TGFβ antibody to this signaling complex, we computationally generated a composite model of the αvβ8-LTGFB1-GARP-TGFBR2 signaling assembly by overlaying experimentally determined structures of subcomplexes (Figure S1A, left). Superimposition and comparison of this assembly with the experimental structures of the anti-TGFβ scFv antibody bound to TGFβ and a structure of a full IgG antibody (Figure S1A, right) suggest that the anti-TGFβ antibody epitope is in close proximity to the plasma membrane (Figure S1B). We therefore hypothesized that the anti-TGFβ antibody would be limited in its ability to block the activation induced by integrin αvβ8 interaction with the TGFβ latent complex (Figure S1C).

In vitro assays designed to test TGFβ activity, either with recombinant active TGFβ1 or induced by integrin αvβ8 activation, allowed us to further investigate the mechanism of action of the anti-TGFβ antibody compared to the anti-αvβ8 antibody. In a reporter cell assay designed to test the ability to block the active TGFβ1 cytokine, the anti-TGFβ antibody was active in the nM range (9.44 nM IC50), while the anti-αvβ8 antibody was, as expected, inactive (Figure 1A). On the other hand, when the assay was designed to test the ability to block integrin αvβ8-dependent TGFβ activation (with releasable TGFβ [WT] or with a mutated non-releasable R249A TGFβ [NR] Zhang et al 2026), using a co-culture cell-based reporter assay in which one cell type express integrin αvβ8 and the other express the TGFβ1-GARP complex and the SMAD reporter cassette, the anti-αvβ8 antibody showed potent inhibition (WT TGFβ1 IC50, 0.17 nM; NR TGFβ1 IC50, 0.089 nM), while the anti-TGFβ antibody was only able to block the pathway at a much higher concentrations (IC50, 79.55 and 20.32 nM, respectively) (Figure 1B).

**Figure 1.**
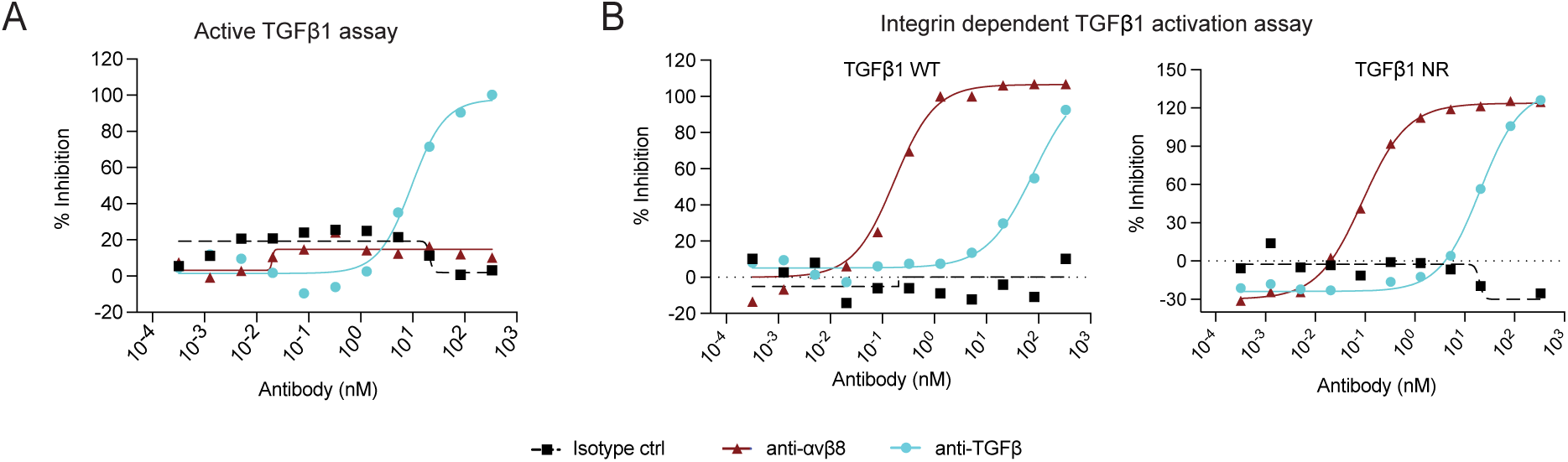
*In vitro* assessment of the TGFβ pathway inhibition ability of anti-TGFβ and anti-αvβ8 antibodies. (A) *In vitro* anti-TGFβ antibody (blue circles) and anti-αvβ8 antibody (purple triangles) activity in an assay devised to test the ability to block active TGFβ1 (representative experiment of many). (B) *In vitro* anti-TGFβ antibody (blue circles) and anti-αvβ8 antibody (purple triangles) activity in an assay devised to test the ability to block integrin-dependent TGFβ1 (wild type, WT, left) or (non-releasable, NR, right) activation (representative experiment of many). IC₅₀ values were determined by nonlinear regression using a four-parameter logistic model.

These *in vitro* experiments confirmed that the anti-TGFβ and anti-αvβ8 antibodies differentially inhibit TGFβ signaling: anti-αvβ8 selectively inhibits signaling induced by αvβ8-mediated activation, and anti-TGFβ preferentially inhibits signaling induced by mature TGFβ.

### Integrin αvβ8 expression is increased in human tumors, and αvβ8 inhibition, when combined with CIT, results in tumor regression in mouse models

We evaluated *ITGB8* (the gene encoding the beta subunit of integrin αvβ8) expression in organ-matched normal and tumor tissue using bulk RNA-seq data from The Cancer Genome Atlas (TCGA) and The Genotype-Tissue Expression (GTeX). *ITGB8* was upregulated in tumors compared to normal tissues in several tissue types, including glioma, glioblastoma, esophageal, lung, kidney, renal papillary, ovarian, and stomach cancer (Figure 2A).

**FIGURE 2.**
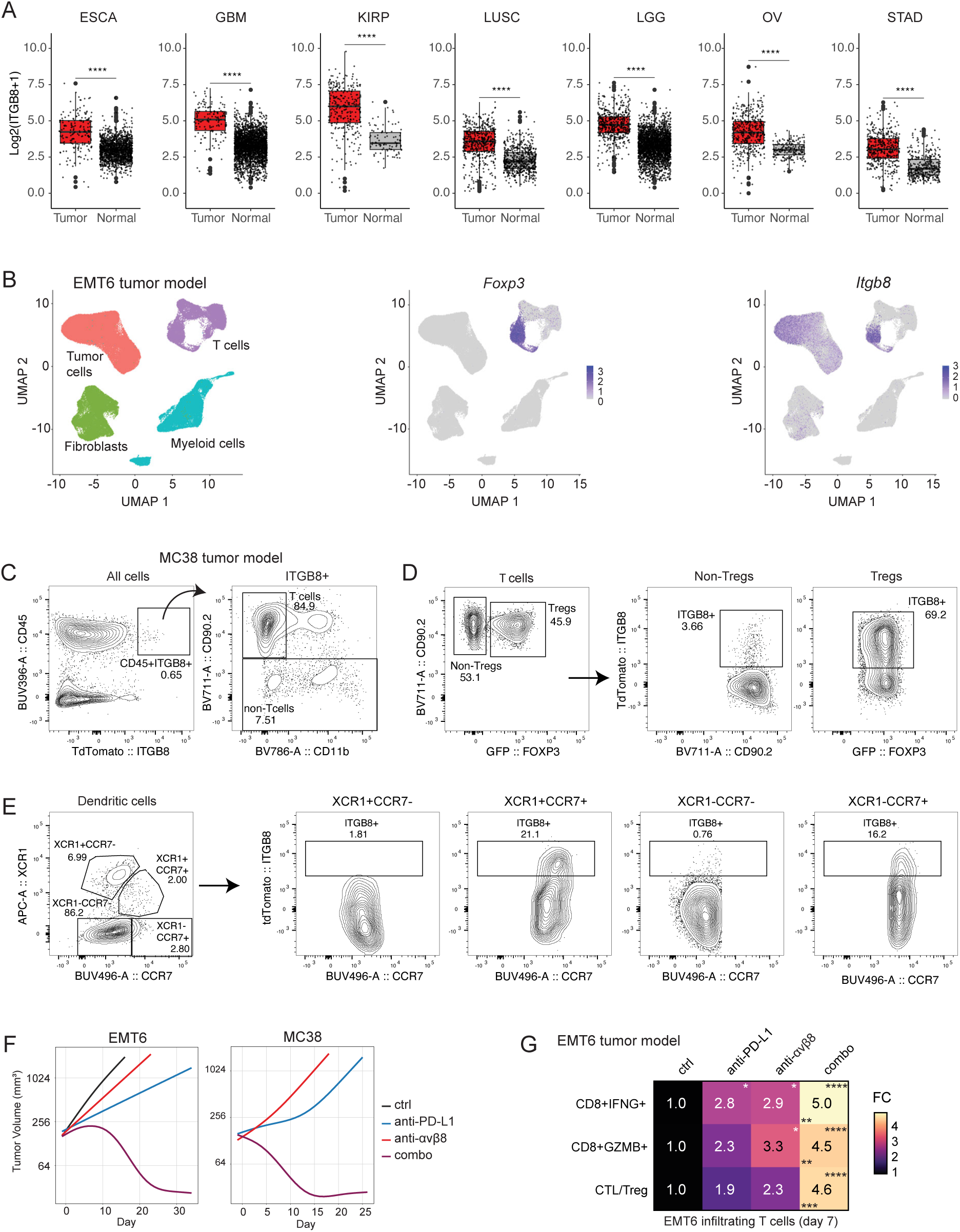
Integrin αvβ8 tumor expression and anti-tumor activity of the anti-αvβ8/anti-PD-L1 combination treatment. (A) ITGB8 expression across tumor indications and the respective normal tissue (TCGA, GTeX). (B) UMAP of cell populations sorted from EMT6 tumors implanted orthotopically in Balbc mice^24^ (left panel). Foxp3 (middle panel) and Itgb8 (right panel) expression levels [Log(CPM/100 + 1)] in UMAP space are shown. (C) Flow cytometry analysis of all TME cells from MC38 tumors implanted in FOXP3-GFP/ITGB8-TdTomato reporter mice (representative of 5 mice). (D) Flow cytometry analysis of T cells from MC38 tumors implanted in FOXP3-GFP/ITGB8-TdTomato reporter mice (representative of 5 mice, gating strategy in Fig S1B). (E) Flow cytometry analysis of DC from MC38 tumors implanted in FOXP3-GFP/ITGB8-TdTomato reporter mice (representative of 5 mice, gating strategy in Fig S1B). (F) Group fit curves of the volume (y-axis) of EMT6 (left panel) and MC38 (right panel) tumors treated with anti-PD-L1 and anti-αvβ8 alone or in combination over time (x-axis; n=10 mice per group, individual curves are shown in Figure S1C). (G) Flow cytometry analysis of EMT6 tumor-infiltrating T cells at 7 days after initiation of treatment (CTL/Treg = CD8+GZMB+ / Treg ratio) (n=9-15 mice per group, from 3 independent experiments, FC = fold change over control group). Tukey’s multiple comparisons test adjusted P values are shown for ctrl vs other treatment (right top corner) and for anti-PD-L1 vs combo a-αvβ8 (bottom left corner of the combo’s cells) *, adj P value <0.05; **, adj P value <0.01; ***, adj P value <0.001; ****, adj P value <0.0001.

We next evaluated αvβ8 expression in syngeneic mouse tumor models. To evaluate integrin αvβ8 expression in the mouse TME, two different approaches were taken. First scRNA-seq analysis of cells sorted from the EMT6 tumor model^24^ showed that both *Foxp3*+ T cells (T_REG_s) and tumor cells expressed *Itgb8* (Figure 2B). Second, the MC38 model TME was evaluated in a *Foxp3*-GFP/*Itgb8*-TdTomato reporter mouse^35^. Flow cytometry analysis confirmed that in the MC38 tumor model, more than 80% of the CD45+ *Itgb8*-TdTomato cells were T cells (Figure 2C), and in particular, that approximately 70% of the T_REG_s expressed *Itgb8*-TdTomato (Figure 2D). Additionally, in-depth characterization of the tumor-infiltrating DCs showed that a fraction of CCR7+ (migratory) DCs expressed *Itgb8*-TdTomato in both XCR1+ (DC1) and XCR1-(DC2) populations (Figures 2E and S2A, B).

As previously shown ^25,28,36^, integrin αvβ8 blockade in combination with a checkpoint inhibitor (anti-PD-L1) induced an efficacious anti-tumor response superior to that of anti-PD-L1 single agent, consistent with prior observations of anti-TGFβ in combination with a checkpoint inhibitor ^5,24^ (Figures 2F and S2C). Also similar to TGFβ ligand blockade^5,24^, the anti-tumor activity induced by anti-αvβ8 in combination with anti-PD-L1 was accompanied by an increased activation of CD8 T cells as shown by the increased percentage of interferon gamma (IFNγ)+ and granzyme B (GZMB)+ CD8 T cells infiltrating the EMT6 TME (Figure 2G). A robust increase in the number of CD8 and CD4 T cells was independently observed in two laboratories (at Genentech, GNE, and at the University of California, San Francisco, UCSF) using two different anti-integrin αvβ8 antibodies combined with either anti-PD-L1 (GNE) or anti-PD-1 (UCSF) (Figure S2D, E). Tumor-infiltrating T_REG_s increased significantly in mice treated with anti-αvβ8 single agent (Figure S2F), but the increase in CD8+GZMB+ T cells (cytotoxic T lymphocytes [CTLs]) was relatively larger, so that in both single agent and in combination with antiPD-L1, the CTL:T_REG_ ratio was higher than in the control group (2.3 and 4.6 folds respectively, Figure 2G and S2F). Both laboratories observed that CD8 T cell activation was significantly augmented in combination treatments, as evidenced by increases in both percentage and mean fluorescence intensity (MFI) of GZMB and IFNγ in CD8 T cells (Figures 2G and S2G, H).

### Anti-αvβ8 and anti-TGFβ antibodies differ in their ability to inhibit the TGFβ pathway in the tdLN

At first glance, anti-tumor activity in mouse tumor models suggested that TGFβ-ligand and integrin αvβ8 inhibition had similar effects. However, TGFβ-blocking antibodies would be expected to inhibit TGFβ activated by integrin-independent mechanisms, whereas anti-αvβ8 antibodies would not. As noted above, recent structural insights^21^ raised the possibility that TGFβ-blocking antibodies might be constrained from inhibiting TGFβ activated by integrin αvβ8. We therefore decided to systematically compare the enhancement of anti-tumor immunity induced by TGFβ and by blocking antibodies targeting integrin αvβ8We next conducted *in vivo* experiments to determine the potency of each antibody in inhibiting TGFβ signaling in the TME and the tdLN. EMT6 or MC38 tumor-bearing mice were treated with an anti-TGFβ or anti-αvβ8 antibody, and TGFβ signaling inhibition was determined by quantifying the percentage of phosphorylated SMAD2/3 (pSMAD2/3) by ELISA. At 24 hours post injection, in both EMT6 tumor and tdLN, there was no significant difference between isotype control and anti-PD-L1 treated animals (Figure S3A), whereas the effects of anti-TGFβ or anti-αvβ8 antibodies in combination with anti-PD-L1 were clearly distinct. Both antibodies were active in the tumor, with anti-αvβ8 showing greater inhibitory activity than the anti-TGFβ antibody (∼50% pSMAD2/3 reduction vs. ∼25%, respectively). In contrast, only anti-αvβ8 was active in the tdLN, with inhibitory activity on par with the assay’s positive control (a small-molecule inhibitor of ALK5[TGFBR1] administered i.v. one hour before tissue collection; Figure 3A). A similar inhibition pattern induced by anti-αvβ8 treatment was observed in the MC38 tumor model and corresponding tdLN (Figure S3B). At 7 days after initiation of treatment, TGFβ pathway inhibition in the tumor was further tested by pSMAD2/3 ELISA (proximal signaling readout) and by analyzing the Fibroblast TGFβ Response Signature (FTBRS) by bulk RNAseq, as readout of downstream TGFβ signaling in the TME ^5^. At this later time point, tumors from mice treated with anti-PD-L1 alone showed a modest reduction in pSMAD2/3 levels compared with controls, suggesting a possible indirect inhibitory effect of checkpoint inhibition on the TGFβ pathway. However, pSMAD2/3 reduction was significantly more pronounced in tumors of mice treated with anti-αvβ8 alone or in combination with anti-PD-L1 (Figure S3C). These results were confirmed by the FTBRS analysis of tumor samples, which showed both single-agent and combinatorial activity of anti-αvβ8 (Figure 3B, right panel). When a similar analysis was run with the anti-TGFβ antibody, FTBRS reduction was observed only in combination with anti-PD-L1, but not with the anti-TGFβ antibody as a single agent^5^ (Figure 3B, left panel).

**FIGURE 3.**
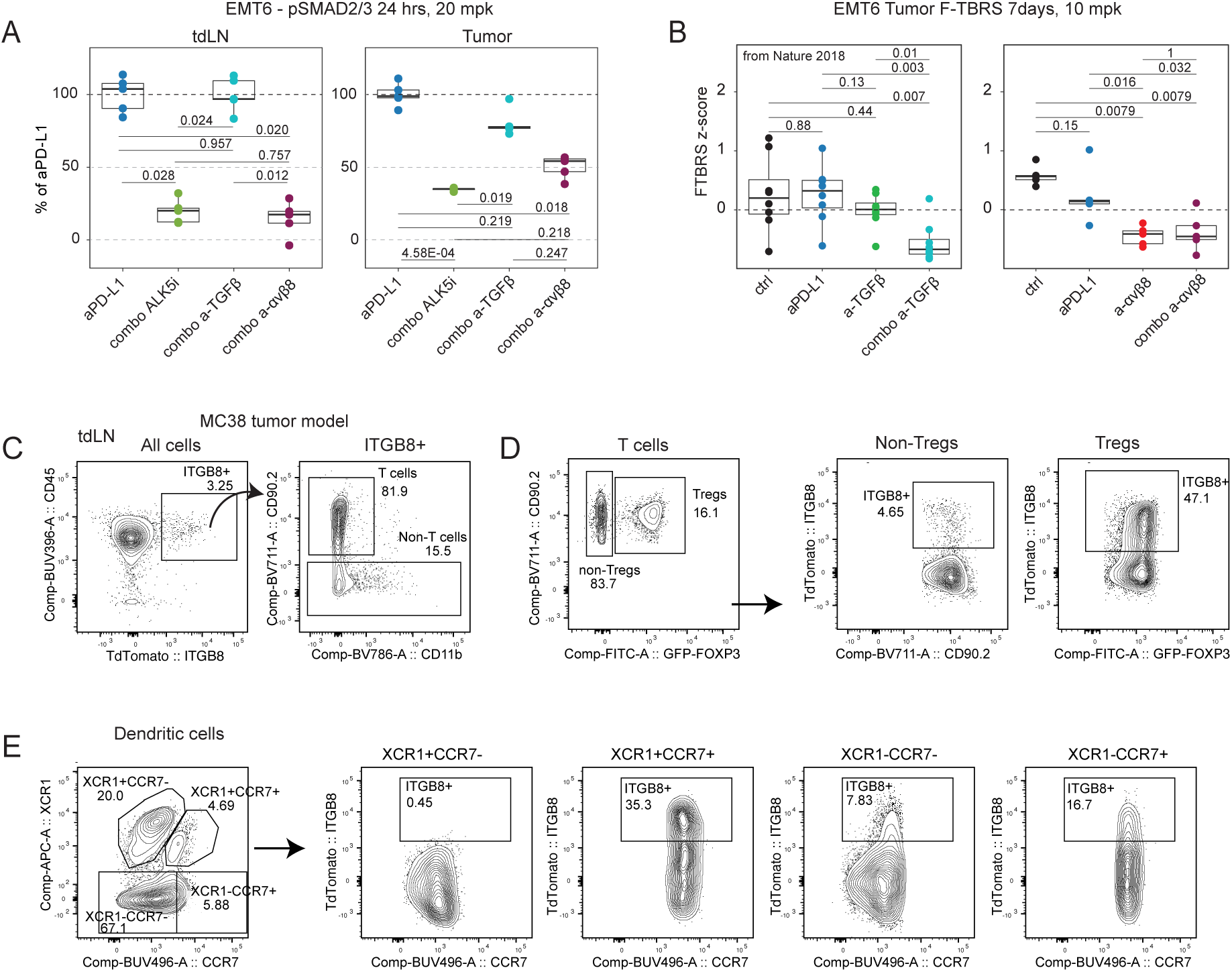
*Ex vivo* assessment of TGFβ pathway inhibition ability of anti-TGFβ and anti-αvβ8 antibodies. (A) *Ex vivo* assessment of SMAD2/3 phosphorylation (pSMAD2/3) by ELISA. Tumor draining lymph nodes (tdLN) and tumors from mice treated with anti-PD-L1 alone or in combination with either an ALK5 small molecule inhibitor (1hr), ant-TGFβ antibody (24 hrs), or anti-αvβ8 antibody (24 hrs) (n=5 mice per group of treatment. Stats: Dunn test with BH correction, adjusted P values are shown). (B) *Ex vivo* RNAseq analysis of tumors from mice treated with anti-TGFβ antibody (left, data from Nature 2018^5^) and anti-αvβ8 antibody (right, n=5 mice per treatment group) at 7 days after initiation of treatment (F-TBRS= fibroblast TGFβ response signature). Stats: Wilcox test with Holm correction; adjusted P-values are shown. (C) Flow cytometry analysis of all cells from the tumor-draining lymph node from MC38 implanted in FOXP3-GFP/ITGB8-TdTomato reporter mice (representative of 5 mice). (D) Flow cytometry analysis of T cells from the tumor-draining lymph node from MC38 tumors implanted in FOXP3-GFP/ITGB8-TdTomato reporter mice (representative of 5 mice). (E) Flow cytometry analysis of DC from the tumor-draining lymph node from MC38 tumors implanted in FOXP3-GFP/ITGB8-TdTomato reporter mice (representative of 5 mice).

The importance of the activity in the tdLN for the anti-tumor response triggered by anti-αvβ8 treatment became more apparent in dose-response studies. The anti-tumor efficacy of anti-αvβ8 in combination with anti-PD-L1 was tested at anti-αvβ8 doses ranging from 0.01 mg/kg to 10 mg/kg in the EMT6 tumor model. All doses showed better anti-tumor activity than the anti-PD-L1 single agent, with maximum activity at 0.1 mg/kg or higher (Figure S3D). In terms of TGFβ pathway inhibition in the tumor, at 24 hours after drug administration, all doses showed some reduction in pSMAD2/3, with maximum inhibition at 1 mg/kg or higher (Figure S3E, right panel). At the same time point, in the tdLN, maximal pSMAD2/3 reduction plateaued at 0.1 mg/kg (Figure S3E, left panel), similar to the dose-response for anti-tumor efficacy.

Given the unique ability of the anti-αvβ8 antibody to reduce TGFβ signaling in the tdLN, αvβ8 expression was analyzed in the tdLN of *Foxp3*-GFP/*Itgb8*-TdTomato reporter mice inoculated with MC38 tumors. 3.25% of lymph node cells were marked by the *Itgb8*-tdTomato reporter and ∼ 80% were T cells (Figure 3C). Half of the T_REG_s in the tdLN expressed *Itgb8*-tdTomato, and approximately 5% of FOXP3-T cells (non-T_REG_s) also expressed *Itgb8*-tdTomato (Figure 3D). ScRNA-seq analysis of T cells in EMT6 tdLNs confirmed that a significant fraction of T_REG_s expressed *Itgb8*, while among CD8 T cells *Itgb8* was enriched in Tpex (*Tcf7*+, *Pdcd1*+, *Lag3*+, *Tox*+, *Havcr2*-CD8 T cells, cluster 0) (Figures S3F, G). Compared to tumors (Figure 2E), a larger fraction of DCs expressed *Itgb8*-tdTomato in the tdLN: 35% of the XCR1+CCR7+ DC, 17% of the XCR1-CCR7+DC, and 8% of the XCR1-CCR7-DC (Figure 3E). This agrees with previous work in naive mice which showed that expression of *Itgb8* is consistently lower in resident than migratory DCs in mesenteric LNs, suggesting that migration and maturation of DCs are required for high expression of αvβ8 ^35,37^.

To summarize, we showed that anti-TGFβ and anti-αvβ8 antibodies differentially affect TGFβ signaling in vivo, consistent with our in vitro observations. Furthermore, these data suggest that integrin αvβ8-mediated TGFβ activation accounts for essentially all TGFβ signaling in the tdLN.

### Anti-tumor effects of αvβ8 inhibition in combination with CIT depend on T cell egress from lymph node, but not on interferons

Because αvβ8 blockade inhibited TGFβ signaling more effectively than TGFβ ligand inhibition in tdLN, we speculated that T cell activation within the lymph node may contribute to the therapeutic activity of αvβ8 inhibition. It has been previously shown that the anti-tumor activity of combined anti-TGFβ and anti-PD-L1 blockade is partially inhibited by blocking the egress of primed T cells from the lymph node (by treatment with FTY-720)^24^. In contrast, we found that FTY-720 completely abolished the activity of combination therapy with anti-αvβ8 and anti-PD-L1, suggesting that mobilization of T cells from tdLN is a mechanistically critical aspect of αvβ8 blockade (Figures 4A-B and S4A).

**FIGURE 4.**
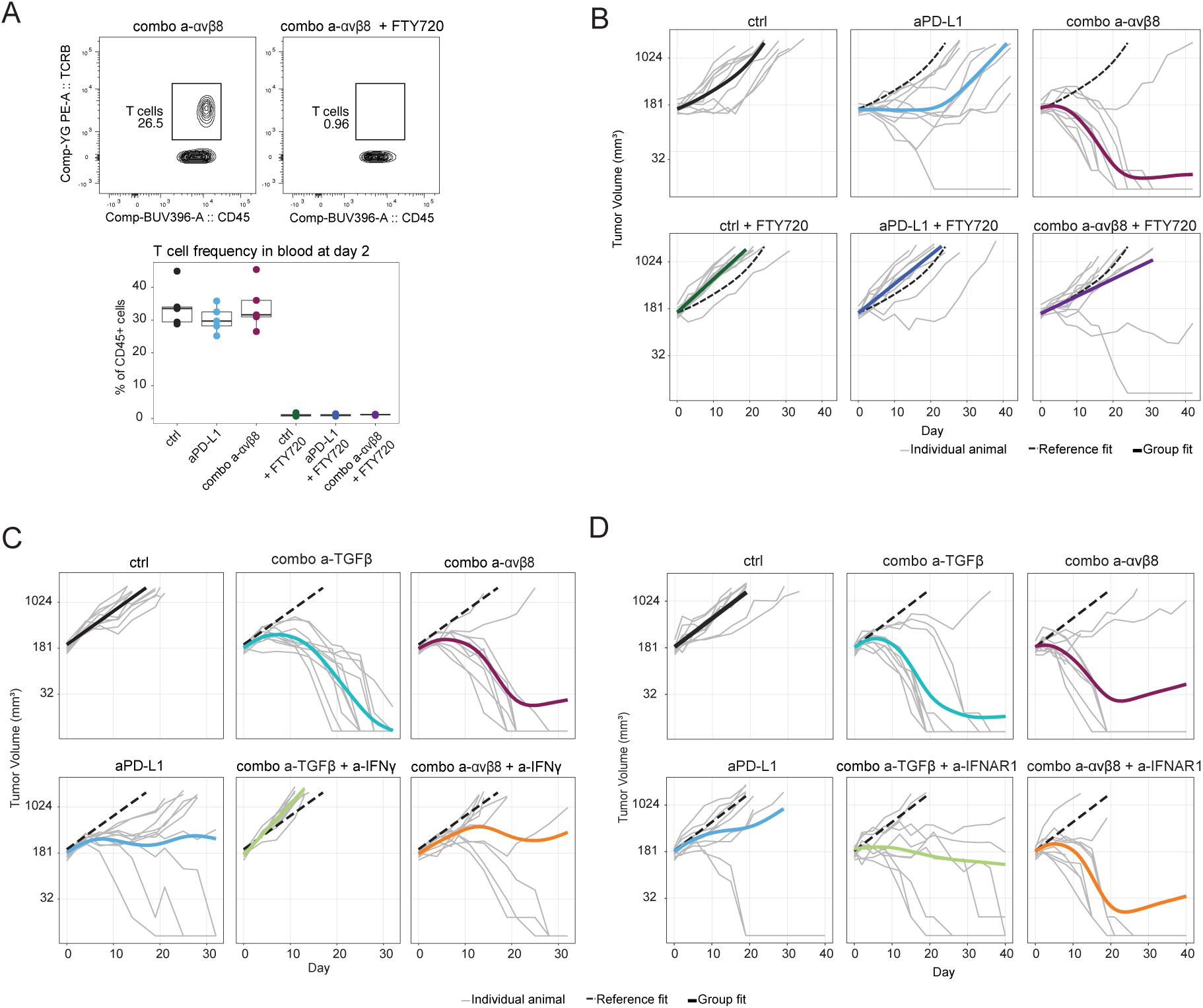
Effects of FTY720 and IFNs inhibition on anti-tumor activity of anti-αvβ8/anti-PD-L1 and anti-TGFβ/anti-PD-L1 combination treatments. (A) Flow cytometry analysis of blood from mice treated with anti-αvβ8/anti-PD-L1 combination with or without FTY720, from the experiment shown in panel B. Representative dot plots showing CD45 and TCRB staining (upper panel); quantification of the T cell frequency across treatments at 2 days after initiation of treatment (bottom panel, n=5 per treatment group). (B) Tumor volume (y axis) of EMT6 tumors treated with anti-αvβ8/anti-PD-L1 combination with or without FTY720 over time (x axis). Individual animal curves (grey lines) and group fit curves (thick solid lines) of the control group (black) and treatment groups (colored) are provided (n = 10 for all groups). (C) Tumor volume (y axis) of EMT6 tumors treated with anti-αvβ8 or anti-TGFβ/anti-PD-L1 or combination with or without anti-IFNγ over time (x axis). Individual animal curves (grey lines) and group fit curves (thick solid lines) of the control group (black) and treatment groups (colored) are provided (n = 10 for all groups). (D)Tumor volume (y axis) of EMT6 tumors treated with anti-PD-L1/ αvβ8 or anti-TGFβ combination with or without anti-IFNAR1 over time (x axis). Individual animal curves (grey lines) and group fit curves (thick solid lines) of the control group (black) and treatment groups (colored) are provided (n = 10 for all groups).

It was also previously demonstrated that the anti-tumor response induced by the anti-TGFβ/anti-PD-L1 combination was dependent on IFNγ^24^, consistent with its documented role in promoting anti-tumor immunity. To test if the response induced by the anti-αvβ8/anti-PD-L1 combination similarly required IFNγ, mice bearing EMT6 tumors were treated with a combination of either anti-TGFβ /anti-PD-L1 antibodies or anti-αvβ8/anti-PD-(L)1 with or without the addition of IFNγ neutralizing antibody. While the anti-tumor efficacy of combined anti-TGFβ /anti-PD-L1 blockade was strictly dependent on IFNγ, anti-αvβ8 in combination with either anti-PD-L1 or anti-PD-1 retained substantial anti-tumor activity in the absence of IFNγ signaling (Figures 4C and S4B). Similarly, when an anti-IFNAR1 antibody was added to neutralize type I interferons, the anti-tumor response induced by the anti-TGFβ/anti-PD-L1 was partially inhibited, while anti-αvβ8/anti-PD-L1 efficacy was not affected (Figure 4D).

Collectively, these data demonstrate that the anti-tumor response induced by αvβ8 inhibition is biologically quite different from the response elicited by TGFβ ligand blockade in mouse models.

### T_REG_s are not differentially affected by anti-αvβ8/anti-PDL1 combo therapy compared with single-agent treatments

As we observed robust integrin αvβ8 expression on T_REG_s in tumor and tdLN, we asked whether the differential effects of the two treatments could be ascribed to distinct effects on T_REG_s. At 7 days after initiation of treatment, T cells were sorted from EMT6 tumors treated with anti-PD-L1, TGFβ or αvβ8 blocking antibodies, or combinations thereof, and analyzed by scRNA-seq. Comparison of T_REG_ gene expression revealed some obvious differences between treatment groups. For example, while expression of *Ccr8* was induced by anti-PD-L1 treatment, it was inhibited significantly by αvβ8 blockade but unaffected by anti-TGFβ (Figure S5A). Similarly, expression of genes known to be important for the immunosuppressive function of T_REG_s (including *Id2, Il2ra, and Tgfb1*), was reduced by αvβ8 blockade but not anti-TGFβ (Figure S5A). However, these changes in T_REG_ gene expression could not, by themselves, account for the anti-tumor efficacy of αvβ8 inhibition, since identical effects were observed with αvβ8 inhibition with or without anti-PD-L1, whereas only combination therapy induced effective anti-tumor immunity (Figure 2F). This conclusion was further supported by a pseudo-bulk analysis of differentially expressed genes (DEGs) in T_REG_s across all four treatments. While anti-αvβ8/anti-PD-L1 combination treatment significantly downregulated expression of immunosuppressive genes (including heat shock proteins that have been shown to be important for immunosuppressive functions of T_REG_s ^38^) compared to control or anti-PD-L1 single-agent treatment, minimal differences were observed between anti-αvβ8 alone or in combination with anti-PD-L1 (Figure S5B).

### Tumor DC transcriptional, phenotypic and functional states are uniquely altered by anti-αvβ8/anti-PD-L1 combination therapy

Next, we turned our attention to DCs, the other major αvβ8-positive cell population detected in both tumor and tdLN. EMT6 tumor-infiltrating cDCs were quantified by flow cytometry as CD45+ Ly6G- F4/80- LY6C- CD11C+ MHCII+ (Figure S6A-B) in mice treated with either anti-TGFβ or anti-αvβ8 antibody alone or in combination with anti-PD-L1. In mice treated with anti-TGFβ /anti-PD-L1 combination treatment, 7 days after initiation of treatment, the number of cDCs was significantly increased compared to control or anti-TGFβ single-agent-treated mice. Anti-PD-L1 single-agent treatment also significantly increased cDCs, while anti-TGFβ single treatment did not (Figures 5A and S6C, left panels). The effect of anti-αvβ8 on the same population of cells was markedly different: while anti-PD-L1 consistently increased tumor-infiltrating cDCs, both anti-αvβ8 single agent and anti-αvβ8/anti-PD-L1 combination treatments significantly decreased the presence of this cell population in the tumor (Figures 5A and S6C, right panels). Further investigation revealed that the cDC reduction in anti-αvβ8 treated mice was driven by a significant loss of CD11b+ XCR1-cells, while cDC1 (defined as CD11b^-^ XCR1^+^ within the cDC gate, Figure S6B) increased with all treatments compared to control (Figure 5B). To better understand the effects of αvβ8 inhibition on dendritic cells phenotypes, scRNA-seq analysis was performed. cDCs (CD45+ CD11b+ Ly6G- F4/80- LY6C- CD11C+ MHCII+) were sorted from EMT6 tumors from mice treated with control, anti-PD-L1, anti-αvβ8, or anti-αvβ8 plus anti-PD-L1 antibodies at 7 days after initiation of treatment. After filtering, 16,803 cells were analyzed, and six clusters were identified based on their gene expression profile (Figure 5C). Three clusters of cDC1s were identified (all expressing *Zbtb46*, *Clec9a, Irf8* and *Xcr1),* and each was characterized by the expression of identifying genes: *Sell* (cluster 0), *Plet1* (cluster 1), and *Cd207* (cluster 2). One cluster of cDC2 cells (cluster 4, expressing *Zbtb46, Clec10a and Sirpa*); one cluster of *Ccr7* expressing migratory DCs (cluster 5); and one cluster of cycling cDC1s (cluster 3, expressing *Mki67* and enriched in cells expressing *Xcr1, Cd207 and Itgae*) were also identified (Figures 5C and S6D-E). Anti-αvβ8 treatments (alone and in combination with anti-PD-L1) significantly changed the distribution of DCs across these clusters (Figure 5D). Migratory DCs (cluster 5) were significantly decreased by αvβ8 inhibition. cDC1 cells (including clusters 0-3) were particularly increased by the combination therapy, while cDC2 cells (cluster 4) were decreased by the combination and anti-PD-L1 single agent compared to control (Figure 5E, top panel). Within the cDC1 populations, *Sell*+ cDC1 (cluster 0) percentages were slightly increased by all treatments, *Plet1*+ cDC1 (cluster 1) were significantly decreased by anti-αvβ8 treatments, while *Cd207*+ (cluster 2) and cycling (cluster 3) cDC1s were increased by anti-αvβ8 treatments (Figures 5E, bottom panel, and S6F).

**FIGURE 5.**
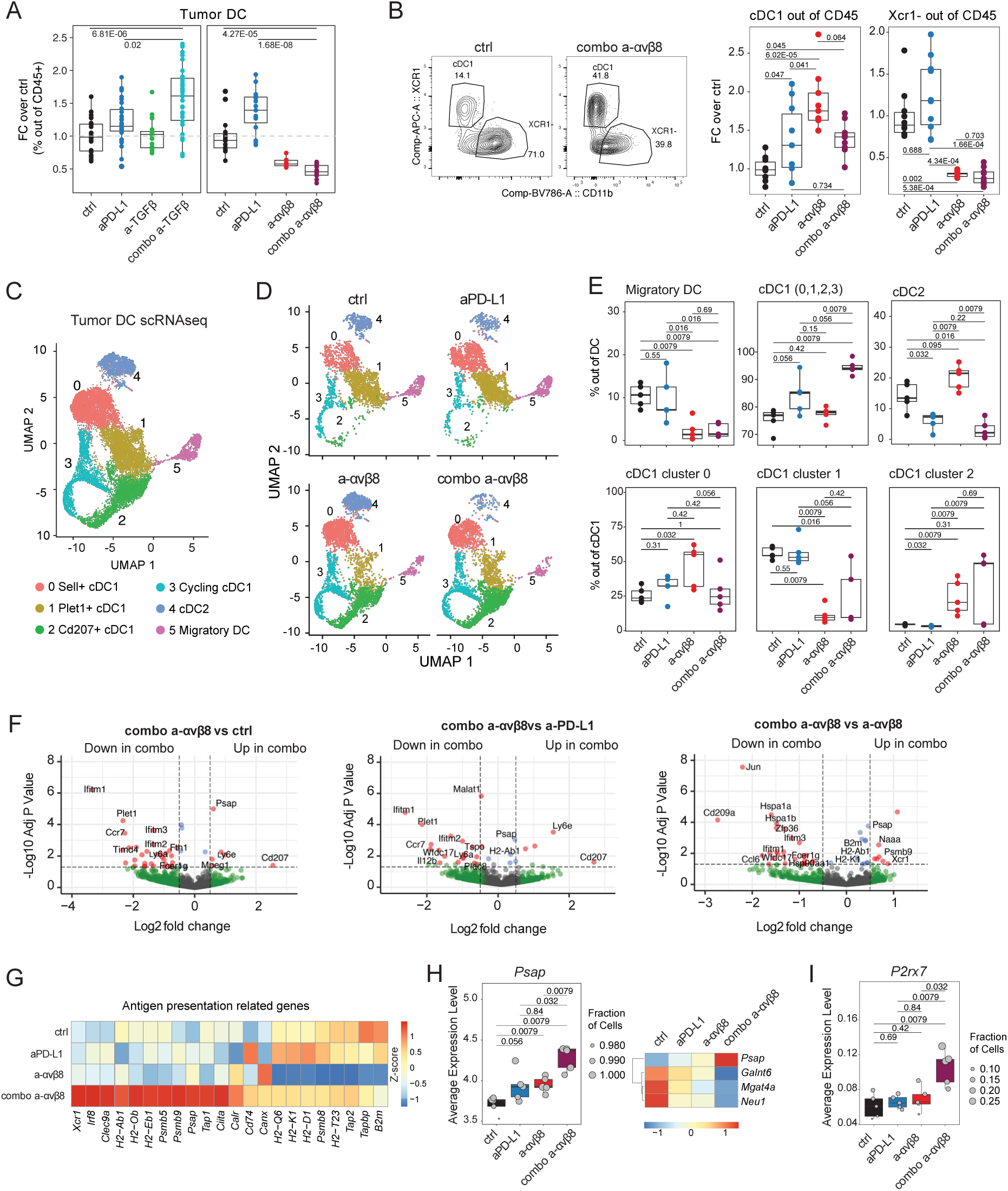
Analysis of tumor DC phenotypes following anti-αvβ8 antibody treatments. (A) Flow cytometry quantification of EMT6 tumor-infiltrating DC within CD45+ cells at seven days after initiation of treatment. Anti- TGFβ / PD-L1 combination (left), anti-αvβ8/anti-PD-L1 antibodies (right). Gating strategy shown in Supplementary figures S5A-B. Fold change over average of the control group (y axis; groups: x axis; data from: six independent anti-TGFβ experiments, ctrl n = 30, aPD-L1 n = 30, a-TGFβ n = 20, combo a-TGFβ n = 30; three independent αvβ8 experiments, ctrl n = 15, aPD-L1 n = 15, a-αvβ8 n = 9, combo a-αvβ8 n = 14). Stats: Dunn test with BH correction, adjusted P values of specific comparisons are shown. (B) Flow cytometry analysis of EMT6 tumor-infiltrating DC at seven days after initiation of treatment. Left panels: representative plots of CD11b (x-axis) and XCR1 (y-axis) staining within the DC population in control and combo a-αvβ8-treated mice. Right panels: quantification of the percentage of cDC1 (XCR1+CD11b-DCs) and XCR1-within CD45+ cells across treatments. Fold change over average of the control group (y axis; groups: x axis; data from two independent αvβ8 experiments, ctrl n = 10, aPD-L1 n = 10, a-αvβ8 n = 9, combo a-αvβ8 n = 9). Stats: Dunn test with BH correction, adjusted P values are shown. (C) UMAP of 16,803 dendritic cells (dots) sorted from EMT6 tumors at seven days after initiation of treatment, colored by cluster (n = 5 mice per group). (D) UMAP of dendritic cells (dots) colored by cluster (same as in C) and separated by group of treatment (n = 5 mice per group). (E) Quantification of the percentage of cells within clusters (y axis; n = 5 per group of treatment from one experiment) in each animal (dots) by treatment group (x axis). The top panel shows percentages within the whole DC object. The bottom panels show percentages within the cDC1 cells. Stats: Wilcox test with Holm correction, adjusted P values are shown. (F) Pseudo-bulk differential expression analysis comparing EMT6 tumor-infiltrating DCs from the anti-αvβ8/anti-PD-L1 combination and either the control (left), the anti-PD-L1 (middle), or the anti-αvβ8 groups (right) (n=5 per group). Dashed lines indicate significance thresholds: fold change, ± 0.5, and adjusted p-value< 0.05. (G) Heatmap of scaled average expression of selected antigen presentation-related genes in DCs for each treatment group. (H) Expression of *Psap* and some of the enzymes involved in its glycosylation in EMT6 tumor infiltrating DC. Left panel: individual animal values are shown with the size of the dots representing the fraction of DCs expressing the specified gene; n=5 per group of treatment; P values are from paired Wilcoxon rank-sum tests (two-sided) comparing between treatment conditions. Right panel: scaled average expression per treatment. (I) Expression of *P2rx7 i*n EMT6 tumor-infiltrating DC (individual animal values are shown with the size of the dots representing the fraction of DCs expressing the specified gene; n=5 per group of treatment; P values are from paired Wilcoxon rank-sum tests (two-sided) comparing between treatment conditions.

To understand how the effects on DC heterogeneity were connected to anti-tumor activity we performed pseudo-bulk analysis of DEGs across treatments in DCs as a whole (Figures 5F and S6G). Some of the DEGs were consistent with the observed changes in cluster frequency. For example, *Ccr7* (a marker of migratory DCs – cluster 5) and *Plet1* (a marker of cluster 1) were significantly downregulated in DCs from combination-treated mice compared to control and anti-PD-L1, while *Cd207* was upregulated in the same comparisons. Similarly, genes expressed by cDC2 (like *Cd209a*) were upregulated in DCs from mice treated with anti-αvβ8 but down-regulated by anti-αvβ8/anti-PD-L1 combination treatment. Interestingly, DEG analysis between anti-αvβ8 single agent and anti-αvβ8/anti-PD-L1 combination treatment highlighted an increase in the expression of genes involved in antigen processing and presentation (enzymes, chaperones, and MHC molecules) including *Xcr1*, *Naaa*, *Psmb9*, *Psap, Tap1, Ciita*, *B2m*, *H2A-b1*, and *H2-K1* (Figures 5F, right panel, and 5G). Interestingly, the expression of *Psap* (prosaposin) was dramatically increased by the anti αvβ8/anti-PD-L1 combination compared to all other treatments (Figure 5H). *Psap* expression was particularly enriched in clusters that increased in response to αvβ8 inhibition (like Cd207+ cDC1 cluster 2 and proliferating cDC1 cluster 3) and lower in clusters that were particularly reduced in combination treatment (cDC2 cluster 4 and migratory DC cluster 5) (Figure S6H). Recently, prosaposin hyperglycosylation has been reported as a TGFβ-dependent tumor resistance mechanism that inactivates DC tumor antigen presentation^39^. The expression of TGFβ-dependent enzymes responsible for prosaposin hyperglycosylation (*Galnt6*, *Mgat4*, *Neu1*) was reduced by all treatments compared to the control (Figure 5H). The combination of higher *Psap* expression with lower expression of its glycosylation would be expected to further enhance prosaposin activity and thus the effectiveness of antigen presentation in mice treated with the anti-αvβ8/anti-PD-L1 combination. Furthermore, the expression of *P2rx7*, an ATP receptor known to be crucial for T cell priming^40,41^, was upregulated in DCs from anti-αvβ8/anti-PD-L1 combination-treated mice (Figure 5I).

To summarize, both anti-TGFβ and anti- αvβ8 in combination with anti-PD-L1 alter tumor-infiltrating DCs. However, these changes differ substantially between the two treatment regimens. In particular, scRNA-seq analysis of tumor DCs sorted from mice treated with anti-αvβ8 revealed significant changes in the composition of the DC populations induced by the therapy, and modulated expression of genes predicted to enhance the antigen-presenting capacity of DCs from mice treated with the anti-αvβ8/anti-PD-L1 combination therapy.

### Lymph node DCs gain enhanced antigen presentation capacity following anti-αvβ8/anti-PD-L1 treatment

The DC clusters identified by scRNA-seq analysis were used to further compare the anti-TGFβ and anti-αvβ8 treatments by flow cytometry. Cells from EMT6 tumors were analyzed at day 7 after treatment initiation. DCs were identified as CD45+ Ly6G- F4/80- LY6C-CD19-CD11C+ MHCII+ and, within this population, CCR7 expression was used to identify migratory DCs (cluster 5). Within the CCR7-XCR1+ cells (cDC1), a combination of CD207 and CD103 expression was used to identify the three cDC1 clusters: CD103^-^CD207^-^ as cluster 0, CD103+CD207-as cluster 1, and CD103+CD207+ as cluster 2 (Figures 6A and S7A). Overall, the scRNA-seq findings relevant to the effects of αvβ8 inhibition were validated by flow cytometry analysis. Comparing antibody treatments confirmed that αvβ8 inhibition affects tumor DCs in a significantly different manner from treatment with TGFβ-blocking antibody, with increases in tumor cDC1s (CCR7-XCR1+CD11b-) induced by αvβ8 inhibition but not by TGFβ-blocking antibody. Migratory DCs (CCR7+ cells) were reduced by all treatments except for anti-PD-L1single agent, and CCR7-XCR1-CD11b+ cells were significantly reduced only by αvβ8/PD-L1 inhibition (Figure 6B, top panel). Among cDC1s, the major differences induced by anti-PD-L1 in combination with either anti-TGFβ or anti-αvβ8 treatments manifest in the fraction of cells in cluster 0 and cluster 2: the anti-TGFβ combination showed a greater increase in the fraction of cells in cluster 0, while the anti-αvβ8 combination increased the fraction of cells in cluster 2 (Figure 6B, bottom panel). Pseudo-bulk analysis of the DEGs between cDC1 was performed. Notably, several genes involved in antigen presentation were upregulated in cluster 2 compared to cluster 0 (Figure 6C). The antigen processing and presentation pathway (GO:0019882) was significantly upregulated in cluster 2 compared to cluster 0 (Figure 6D), suggesting that DCs from mice treated with anti-αvβ8/anti-PD-L1combination treatment might be better at antigen presentation than DCs from mice treated with anti-TGFβ/anti-PD-L1 combination treatment.

**FIGURE 6.**
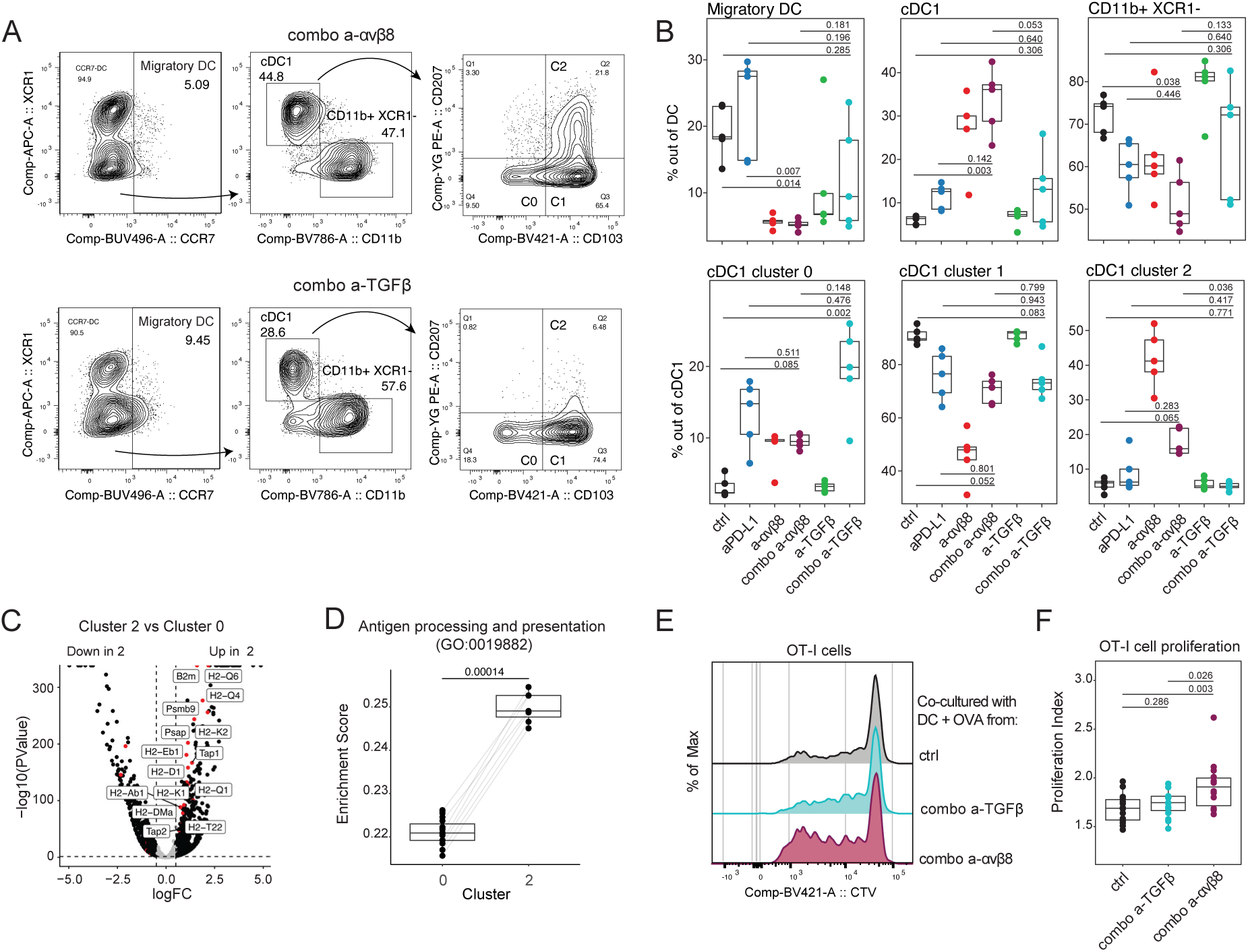
Direct comparison of the effects of anti-αvβ8/anti-PD-L1 and anti-TGFβ /PD-L1 on tumor DCs phenotypes and LN DCs antigen presentation capability. (A) Flow cytometry quantification of the EMT6 tumor-infiltrating DC using the DC clusters identified by scRNAseq. Representative plots with the gating strategy are shown from anti-αvβ8/anti-PD-L1 antibodies (top), and anti-TGFβ/anti-PD-L1 combination (bottom) (upstream gating strategy and CCR7 FMO controls in supplementary Figure S6A). (B) Quantification of the percentage of cells belonging to the different DC populations (y axis; n = 5 per group of treatment from one experiment) in each animal (dots) by treatment group (x axis). Stats: Dunn test with BH correction, adjusted P values are shown. (C) Pseudo-bulk differential expression analysis comparing cluster 0 and cluster 2 cells from UMAP in Figure 4C. Dashed lines indicate significance thresholds: fold change, ± 0.5, and adjusted p-value< 0.05 (DEseq2). Highlighted in red are genes involved in antigen presentation (GO:0019882). (D) Enrichment score for expression of genes from the antigen processing and presentation pathway (GO:0019882) in DCs from clusters 0 and 2. Each dot represents one mouse (n=20). Lines connect scores from the same mouse. (E) Flow cytometry analysis of OT-I cell proliferation based on the cell trace violet (CTV) fluorescence intensity. Representative plots of OT-I cells co-cultured for three days with OVA-pulsed DC from the lymph nodes of mice treated with control, anti-TGFβ / PD-L1, and anti-αvβ8/anti-PD-L1 combination. No OVA controls are shown in Figure S6B. (F) Quantification of OT-I cell proliferation by proliferation index (y axis; n = 14-15 per group of treatment from two experiments) in each animal (dots) by treatment group (x axis). Stats: Anova multiple comparisons; adjusted P-values are shown.

To directly test this hypothesis, DCs from the LN of naive mice treated with either therapy were assessed for their ability to activate OT-I cells. C57Bl6 mice were injected i.v. with either anti-TGFβ/anti-PD-L1 or anti-αvβ8/anti-PD-L1 combination. 24 hours later, DCs were sorted from lymph nodes, pulsed with ovalbumin (OVA), and co-cultured with CD8 T cells from OT-I mice for 3 days. OT-I cells co-cultured with DCs from mice treated with the anti-αvβ8/anti-PD-L1 combination had a significantly higher proliferation index than the same cells cultured with DCs from the control or anti-TGFβ/anti-PD-L1 combination-treated mice (Figures 6E, F; without OVA controls are shown in Figure S6B). These comparative studies suggest that blocking αvβ8-mediated activation of TGFβ, in combination with checkpoint blockade, is distinct in its capacity to increase antigen presentation and T cell activation by DCs compared with neutralizing the active form of TGFβ.

## Discussion

Although TGFβ pathway inhibition in combination with checkpoint inhibitors has been reproducibly effective in preclinical tumor models^2,4,5,42^, the translatability to clinical trials in cancer patients has been underwhelming. Here, we show that not all TGFβ-targeting therapies are equal. Given recent data suggesting that integrin αvβ8 activates TGFβ1/3 without releasing the active cytokine from the latent complex^23^, we focused on comparing the biological effects of antibody treatments that aim to block either the active cytokines from signaling via TGFBR (the major approach tested so far in the clinic) or TGFβ signaling enabled by αvβ8 integrin.

At first glance, the two strategies appeared to be functionally similar. In EMT6 and MC38 tumor models, both αvβ8- and active-TGFβ blocking antibodies combined with an immune checkpoint inhibitor to induce superior CD8 T cell responses and tumor regression relative to anti-PD(L)1 single agent therapy. However, further experimental investigation revealed that the underlying biological mechanisms were clearly distinct.

First, when the two major sites of the cancer immunity cycle^43^ were analyzed, we unexpectedly observed that although both antibodies inhibited TGFβ activity in the TME, only anti-αvβ8 blocked TGFβ activity in the tdLNs. Treatment with FTY720 confirmed that activity of anti-αvβ8 in the tdLN is likely more relevant for the anti-tumor activity of the anti-αvβ8/anti-PD-L1combination treatment, than for the anti-TGFβ/anti-PD-L1 regimen (as previously shown^24^), since prevention of T cell egress from lymph nodes more strongly inhibited the efficacy of the αvβ8 combination treatment. Second, we observed different degrees of interferon dependency: the anti-TGFβ/anti-PD-L1 combination was completely dependent on IFNγ and partially dependent on type I interferon signaling, whereas the anti-αvβ8/anti-PD-L1 combination treatment effect was largely independent of interferon signaling.

Finally, we observed differences at the cellular level. T_REG_s appeared to be less immunosuppressive upon treatment with anti-αvβ8/anti-PD-L1 than with the anti-TGFβ/anti-PD-L1 combination, as indicated by gene expression. However, the number of T_REG_s infiltrating the tumor was not decreased by either combination therapy^5^ (Figure S2F). We found even more dramatic differences in the effects of the two therapies on DCs. The number of tumor-infiltrating DCs was increased by combination therapy with anti-TGFβ and decreased by combination treatment with anti-αvβ8. The reduction in DC numbers in anti-αvβ8-treated tumors appeared to be driven by a decrease in cDC2 cells. In contrast, within tumor-infiltrating DCs, the percentage of cDC1, considered fundamental to anti-tumor immunity^44–46^, was significantly increased. More importantly, in vivo treatment with the anti-αvβ8/anti-PD-L1 combination treatment altered expression of genes in DCs in ways predicted to enhance antigen presentation and increased bona fide antigen presentation by lymph node DCs, whereas treatment with the anti-TGFβ/anti-PD-L1combination had neither effect. An outstanding question not addressed by these studies is whether these effects result from direct, cell-autonomous αvβ8 inhibition of DCs or from indirect effects on DCs, for example, induced by αvβ8 inhibition of T_REG_s or tumor cells.

Our data show that while in the tdLN, αvβ8-dependent activation of TGFβ is the dominant mechanism through which the TGFβ pathway is activated, both αvβ8-dependent and αvβ8-independent activation mechanisms appear to be important in the TME, as both anti-TGFβ and anti-αvβ8 antibodies partially inhibited TGFβ signaling in the tumor, but neither reached the same inhibitory activity as the ALK5 (TGFBR1) inhibitor. As mentioned above, in the TME, the number of DCs, regardless of their phenotype, was increased by combination therapy with the anti-TGFβ antibody. It is possible that this local increase in DC presence results in overall better *in situ* antigen presentation than in control or anti-PD-L1-treated mice, potentially explaining the independence of the anti-tumor response induced by the anti-TGFβ/anti-PD-L1 combination from tdLN activity. Based on our observations, we speculate that αvβ8 inhibition in combination with checkpoint blockade promotes DC migration to the tdLN while simultaneously enhancing their antigen-presenting capacity, making the tdLN the primary site of action for the anti-αvβ8 combination with checkpoint inhibitors. The remaining tumor-infiltrating DCs are also potentiated in their antigen-presenting ability, suggesting that even if LN activity is necessary for anti-tumor activity, some *in situ* antigen presentation still occurs at the tumor site. On the other hand, neutralizing the active TGFβ ligands potentiates only local activity of tumor-infiltrating immune cells induced by anti-PD-L1 therapy within the tumor. This raises the possibility that failure of TGFβ inhibitors tested to date to effectively augment immunotherapeutic benefit in clinical trials of solid tumors may stem from their inability to disable functionally critical pools of TGFβ that are activated in specific tissue and cellular contexts by integrin signaling.

Taken together, these data demonstrate that αvβ8 inhibition represents a differentiated approach to TGFβ pathway blockade. The two approaches for inhibiting TGFβ signaling assessed here — direct targeting of active TGFβ ligands and blocking TGFβ1/3 activation via inhibition of the αvβ8 integrin — result in distinct biological effects and should not be considered equivalent or interchangeable. In light of these substantial mechanistic differences, the modest clinical efficacy observed with prior efforts to enhance anti-tumor immunity by inhibiting TGFβ ligands should not discourage ongoing efforts to enhance anti-tumor immunity in patients by inhibiting the αvβ8 integrin.

## Material and methods

### Mice

Female BALB/c or C57BL6 mice were obtained from Charles River Laboratories (Hollister, CA). All mice housed at Genentech are kept in individually ventilated cages within animal rooms maintained on a 14:10-hour light:dark cycle. Mice housed at UCSF were maintained in a barrier facility. Animal rooms were temperature and humidity-controlled, between 68-79°F and 30-70% respectively, with 10 to 15 room air exchanges per hour. Mice were acclimated to study conditions for at least 3 days before tumor cell implantation. Animals were 8-10 weeks old. Only animals that appeared healthy and free of obvious abnormalities were used in the studies. Animals were maintained in accordance with the *Guide for the Care and Use of Laboratory Animals*. Genentech and UCSF are AAALAC-accredited facilities and all animal activities in the research studies were conducted under protocols approved by either the UCSF or the Genentech Institutional Animal Care and Use Committee (IACUC).

Foxp3^IRES-eGFP^ x *Itgb8*-IRES-tdTomato (*Foxp3*-GFP/*Itgb8*-TdTomato) reporter mice^35^ were maintained at UCSF in accordance with protocols approved by the University of California, San Francisco IACUC.

### Animal studies

EMT6 and MC38 tumor growth experiments were conducted as previously described^5,24^. All antibodies (isotype control: mouse IgG1 or IgG2a anti-gp120; anti–PD-L1: mouse IgG1 clone 6E11; anti PD-1: clone RMP-14, BioXcell #BE0146; anti-TGFβ: mouse IgG1 (GNE); a-αvβ8; mouse IgG2a-LALAPG (GNE clone A) or ADWA-11 (UCSF)) were dosed at 10 mg/kg, unless otherwise specified, 2 times a week or once a week (ADWA11) for 21 days, with the first dose administered intravenously and subsequent doses administered intraperitoneally. For efficacy studies, tumors were measured twice weekly using calipers. For FTY720 experiments, FTY720 was administered orally concurrently with the antibody treatments. FTY720 1mg/kg was administered by oral gavage daily for 21 days.

For IFNγ neutralizing experiments conducted at GNE (Figure 3C), an anti-mouse IFNγ antibody (InVivoPlus Clone XMG1.2, #BP0055; BioXcell) was administered at 12.5mg/kg at the same time as the other antibodies (2 times a week for 21 days, administered intraperitoneally).

For IFNγ neutralizing experiment conducted at UCSF (Figure S3B), an anti-mouse IFNγ antibody (InVivoPlus Clone XMG1.2, #BP0055; BioXcell) was administered intraperitoneally at 10 mg/kg for the first dose and then every three days at 5 mg/kg.

For IFNAR1 blocking experiments (Figure 3D), an anti-IFNAR1 antibody (InVivoPlus Clone MAR1-5A3, #BP0241; BioXcell) was administered at 12.5mg/kg at the same time as the other antibodies (2 times a week for 21 days, administered intraperitoneally).

Mice were euthanized immediately if tumor volume exceeded 2000mm^3^ (maximal tumor volume permitted by the Genentech and UCSF IACUC), or if tumors ever fell outside the IACUC Guidelines for Tumors in Rodents. No mice met the criteria for euthanasia because of body weight loss, nor exhibited adverse clinical signs during the studies.

For flow cytometry analysis or FACS sorting (for subsequent RNAseq analyses or antigen presentation experiments), mice were euthanized seven days after treatment initiation, and tumors were collected.

### Structure composite

The PyMOL Molecular Graphics System v3.1.3 (Schrödinger, LLC) was used to generate a composite model of the αvβ8-LTGFB1-GARP-TGFBR2 signaling assembly by overlaying experimentally determined structures of subcomplexes using the align command. Protein Data Bank identifier (PDB ID) of the subcomplex structures are as follows: aVb8+LTGFB1 (PDB ID 6UJA); GARP+LTGFB1, removing MHG-8 Fab (PDB ID 6GFF) and TGFBR2+TGFB1, removing TGFBR1 and one of the TGFBR2 chains (PDB ID 3KFD). Based on visual inspection, some conformational rearrangements are needed to form this complex without causing steric clashes, but the proposed assembly appears structurally plausible. Using a similar approach, we overlaid the structure of a full IgG antibody, exemplified by the structure of Pembrolizumab (PDB ID 5DK3), with the structure of the anti-TGFβ scFv fragment 1D11 bound to TGFβ (PDB ID4KV5). The proximity of the membrane bilayer to the membrane-anchored TGFBR2 and GARP proteins was assessed by inserting full-length (i.e. containing the transmembrane anchor) Alphafold models of TGFBR2 and GARP (PMID: 37933859) in model membranes using the Positioning of Proteins in Membranes (PPM) Web Server (PMID: 34716622) and comparing these to the composite model of the αvβ8-LTGFB1-GARP-TGFBR2 signaling assembly. This allowed us to draw representative membrane bilayer planes that would be compatible with formation of the signaling assembly and simultaneous membrane anchoring of TGFBR2 and GARP.

### In Vitro Functional Assays

All in vitro assays were performed as previously described^47^. Brief summaries are provided below. For both assays, luciferase activities were measured using the Nano-Glo® Dual-Luciferase® Reporter Assay System (Promega) according to the manufacturer’s instructions. NanoLuc® activity was normalized to firefly luciferase to control for variation in cell number and viability, and IC₅₀ values were determined by nonlinear regression using a four-parameter logistic model.

#### TGFβ1 Inhibition Reporter Assay

TGFβ1 neutralizing activity was evaluated using an NIH 3T3-based dual-luciferase reporter assay as previously described^47^.Briefly, antibodies were prepared as an 11-point, 1:4 serial dilution starting at 333.3 nM and pre-incubated with 20 pM recombinant human TGFβ1 for approximately 1 hour prior to addition to reporter cells. Following incubation for approximately 18 - 20 hours at 37°C in 5% CO₂, luminescence was quantified as described above.

#### αvβ8-Mediated Latent TGFβ Activation Co-culture Assay

Inhibition of αvβ8-mediated latent TGFβ activation was assessed using a previously described dual-luciferase co-culture assay^47^.NIH 3T3-based dual-luciferase reporter cells constitutively expressing GARP-presented latent TGFβ1 (L-TGFβ1 wild type or non-releasable, R249A mutation) were used in this assay. LN-229 cells endogenously expressing αvβ8 were seeded into assay plates and incubated with diluted antibodies (prepared as described above) for approximately 1 hour at 37°C in 5% CO₂. Reporter cells were subsequently added, and co-cultures were maintained for approximately 18 - 20 hours under the same conditions prior to luminescence quantification as described above.

### PSMAD2/3 ELISA

Tumors and lymph nodes were collected at 24 hours or 7 days after antibody treatment started. Tissues were snap frozen in liquid nitrogen upon collection and preserved at -80° C before processing. Samples were placed in an OMNIruptor bead tube with 400 μL of Lysis buffer CST 2X (Cell Signaling Technology #9803), prepared according to the protocol: diluted in H2O and added protease inhibitors (Roche 05 892 791 001) and phosphatase inhibitors (Roche 04 906 837 001), on ice. Samples were then homogenized using a OMNIruptor machine (4m/s; 2×25s, 10s dwell time). Samples were then spun at 16000g for 5 minutes at 4° C, and the proteins in the supernatant were quantified by BCA (Pierce BCA protein assay kit, Thermo Scientific 23227). For the ELISA assays, samples were diluted with the ELISA kit dilution buffer to 5 mg/ml. pSMAD and total SMAD ELISAs were performed following Kit instructions (Total SMAD2/3 ELISA: Cell Signaling Tecnhology #12000 C; pSMAD2/3 ELISA: Cell Signaling Tecnhology #12001). OD ratios were calculated by dividing the OD values of the pSMAD2/3 plate by the OD values of total SMAD2/3 for each sample, and percentages were calculated based on the control samples’ values.

### Preparation of single-cell suspension and antibody staining for flow cytometry analyses and sorting

#### GNE procedures

Tumors and tumor-draining lymph nodes were collected 7 days after treatment initiation. Tissues were weighed and enzymatically digested using a cocktail of dispase, collagenase P, and DNaseI for 45 min at 37°C, to obtain a single-cell suspension. Cells were counted using a Vi-CELL XR (Beckman Coulter, Brea, CA).

For flow cytometry analysis, cells were first incubated with mouse BD Fc block (5 µg/ml, cat. n. 553141) and LIVE/DEAD® Aqua Fixable Dead Cell Stain (Invitrogen, cat. n. L34957) for 30 min on ice. The cells were then stained with fluorochrome-conjugated antibodies (see table). Flow Cytometry data were collected with a BD FAC Symphony flow cytometer (BD Biosciences, San Jose, CA) using BD FACS Diva software (versions 8.0.1 and 9.1, BD Biosciences, San Jose, CA). Flow data were analyzed in FlowJo, exported to CSV files, and statistical analysis was performed using the R package tidyverse, dplyr, and FSA. Treatment-induced changes were identified using a Kruskal-Wallis ANOVA followed by Dunn’s test, with multiple testing correction via the Benjamini-Hochberg method.

For sorting (scRNA-seq), cells were first enriched for live cells using a Dead Cell Removal Kit following the manufacturer’s instructions (Miltenyi Biotec, cat. n. 130-090-101). Cells were then incubated with mouse BD Fc block (5 µg/ml, cat. n. 553141) for 30 min on ice. And then stained for 30 minutes on ice (see table for antibody used), including 5 hashtags antibodies to identify the mice in each treatment group (C0301-5, Biolegend). Just before sorting, cells from the same treatment group were pulled together and stained with 7AAD and eBioscience™ Calcein Blue AM Viability Dye. Cells were sorted using a BD FACSAria™ Fusion flow cytometer (BD Biosciences, San Jose, CA) in 300 μL of MACS buffer kept at 4° C. Cells were counted and resuspended to an appropriate concentration for downstream experiments.

#### UCSF procedures

The excised tumors were placed in a Petri dish with C10 media (RPMI 1640, HEPES 1%, Penicillin/Streptomycin 1X, fetal calf serum 10%, sodium pyruvate 1 mM, non-essential amino acids 1X, and beta-mercaptoethanol 0.45%) and minced with sterile razor blades. The tissue slurry was then transferred to a 50 mL conical tube and mixed with a digestion cocktail containing Collagenase XI (Sigma C9407) at 2 mg/mL, Hyaluronidase (Sigma H3506) at 0.5 mg/mL, and DNase (Sigma DN25) at 0.1 mg/mL, prepared in C10 medium. Cells were incubated in a shaker at 255 rpm for 30 minutes at 37° C. After incubation,15 mL of C10 medium was added to the digestion slurry and placed on ice. The cell slurry was passed through a 100 μm mesh strainer (Falcon #352360) into a clean 50 mL conical tube. Cells were pelleted by centrifugation for 5 minutes at 200 x g at 4° C and mixed with RBC lysis buffer (Sigma R7757). The cell RBC mixture was incubated at room temperature for 5 minutes, then passed through a 40 μm mesh strainer (Falcon #352340) and washed with 1X PBS. The single cell suspension was centrifuged as before and resuspended in 1X PBS. Cell counts were performed using a hemocytometer (Fisher Scientific #02-671-6). Single-cell suspensions were used for subsequent cell surface and intracellular staining. Samples were counted, and approximately 10^6^-10^7^ cells/well were transferred to a V-shaped 96-well plate. Fc receptor and non-specific binding was blocked with anti-CD16/30 (eBioscience #14061) for 10 minutes at 4° C. For intracellular flow cytometry, cells were incubated with 1:1000 eFluor 780 Fixable viability dye (eBioscience #65-0865-14) for 20 minutes at 4° C. Cells were surface stained for 25 minutes at 4 C° and fixed with 4% PFA for 20 minutes at 4° C. The suspension was transferred to 1X Perm buffer (eBioscience #88-8824) for 20 minutes at room temperature, followed by intracellular staining for 20 minutes at 4° C. After staining, cells were transferred into flow cytometry buffer (PBS with 2%FBS, Penicillin/Streptomycin/Glutamate, EDTA 2 mM) for analysis. Prior to intracellular cytokine staining, cell suspensions were stimulated. The cell suspension was counted, and 3 × 10^6^ cells/well were seeded in a round-bottom 96-well plate. Cells were incubated in 200 μL stimulation cocktail (Inomycin, PMA, Brefeldin-A, and Monensin 500x stimulation cocktail (Tonbo #TNB4975-μL 100)) diluted in C10 media for 5 hours at 37° C in 5% CO2.

##### Flow cytometry antibodies

*Flow cytometry antibodies:*
| COLOR | MARKER | CLONE | SOURCE | CAT N |
| --- | --- | --- | --- | --- |
| APC | XCR1 | ZET | Biolegend | 148206 |
| APCCy7 | Ly6G | 127624 | Biolegend | 127624 |
| APCCy7 | CD19 | 6D5 | Biolegend | 115530 |
| APCCy7 | F4/80 | BM8 | Biolegend | 123118 |
| APCCy7 | Ly6C | HK1.4 | Biolegend | 128026 |
| BV421 | CD103 | 2E7 | Biolegend | 121421 |
| BV605 | CD11c | N418 | BD biosciences | 563057 |
| BV711 | CD90.2 | 30-H12 | Biolegend | 105349 |
| BV785 | CD11b | M1/70 | Biolegend | 101243 |
| BUV495 | CCR7 | 4B12 | BD biosciences | 741195 |
| BUV395 | CD45 | 30-F11 | BD biosciences | 564279 |
| BUV737 | MHCII | AF6-120.1 | BD biosciences | 569176 |
| PE | CD207 | 4C7 | Biolegend | 144204 |
| APC | FOXP3 | FJK-16s | invitrogen | 17-5773-82 |
| APCCy7 | CD8a | 53-6.7 | Biolegend | 100714 |
| BV605 | CD45 | 30-F11 | Biolegend | 103140 |
| BV711 | CD4 | RM4-5 | Biolegend | 100557 |
| FITC | GZMB | NGZB | Invitrogen | 11-8898-82 |
| PE | TCRb | H57-597 | Biolegend | 109208 |
| PeCy7 | IFNg | XMG1.2 | Biolegend | 505826 |
| buv395 | CD25 | PC61 | BD Horizon | 564022 |
| BV650 | TER-119 | TER-119 | Biolegend | 116235 |
| APC | TCRb | H57-597 | Biolegend | 109212 |
| BV711 | F4/80 | BM8 | Biolegend | 123108 |
| BV785 | CD11b | M1/70 | Biolegend | 101206 |
| Alexa488 | Thy1.1 | OX-7 | Biolegend | 202506 |
| FITC | Ly6G | 1AB-Ly6g | Invitrogen | 11-9668-82 |
| FITC | Nk1.1 | S17016D | Biolegend | 156507 |
| FITC | CD19 | 6D5 | Biolegend | 115505 |
| PerCp Cy5.5 | Ly6C | HK1.4 | Invitrogen | 45-5932-82 |
| PeCy7 | MHCII | M5/114.15.2 | Biolegend | 107630 |
| APC | CD3 | 17A2 | BioLegend | 100236 |
| PE | GZMB | NGZB | Invitrogen | 12889882 |
| PE/Cy7 | CD8a | 53-6.7 | BioLegend | 100722 |
| FITC | CD4 | RM4-5 | BioLegend | 100510 |
| BV421 | CD45 | 30-F11 | BioLegend | 103134 |
| BV650 | IFNg | XMG1.2 | BioLegend | 505832 |

#### scRNA-seq sample processing

After FACS sorting, cells were processed using the Chromium Single Cell Gene Expression 5′ (V3: avb8 DC dataset in figures 4, 5 and S5, or V2: untreated LN dataset in Figure S2) Library and Gel Bead kit following the manufacturer’s instructions (10x Genomics). Cells were counted, assessed for viability, and then injected into microfluidic chips to form Gel Beads-in-Emulsion (GEMs) in the 10x Chromium instrument. Reverse transcription was performed on the GEMs, and reverse-transcribed products were purified and amplified. Gene expression libraries and hashtag libraries were made from the cDNA, profiled using a Bioanalyzer High Sensitivity DNA kit (Agilent Technologies), and quantified with a Kapa Library Quantification kit (Kapa Biosystems). HiSeq 4000 (Illumina) was used to sequence the libraries.

#### Bulk RNAseq Tissue prep

For Bulk RNAseq tumor samples were collected at 7 days after initiation of therapy and flash frozen in liquid nitrogen. RNA was extracted fowling the protocol of RNeasy mini kit - QIAGEN (74104).

#### Computational analyses

##### Bulk RNA-seq analysis of human data from TCGA and GTeX

GTeX data was taken from https://storage.googleapis.com/gtex_analysis_v8/rna_seq_data/GTEx_Analysis_2017-06-05_v8_RNASeQCv1.1.9_gene_tpm.gct.gz. TCGA data was obtained from https://xenabrowser.net/datapages/?dataset=tcga_RSEM_gene_tpm&host=https%3A%2F%2Ftoil.xenahubs.net&removeHub=https%3A%2F%2Fxena.treehouse.gi.ucsc.edu%3A443. TPMs were log2 normalized with a pseudocount of 1. Only primary tumor and normal samples were kept.

##### Bulk RNA-seq analysis of mouse data

Bulk RNA-seq data were processed using the BioConductor package HTSeqGenie (version 4.4.2, (https://bioconductor.org/packages/ release/bioc/html/HTSeqGenie.html) as follows: first, reads with low nucleotide qualities (70% of bases with quality <23) or matches to rRNA and adapter sequences were removed. The remaining reads were aligned to the human reference genome (GRCm38.p5) using GSNAP (25) version “2013–11–01,” allowing maximum of two mis-matches per 75 base sequence (parameters: -M 2 -n 10 -B 2 -i 1 -N 1 -w 200000 -E 1 –pairmax-rna 1⁄4 2000000). Transcript annotation was based on the Gencode genes data base (GENCODE M15). To quantify gene expression levels, the number of reads mapping unambiguously to the exons of each gene was calculated. The F-TBRS score was calculated by taking the average of the z-score of the genes making up the score (Acta2, Actg2, Adam12, Adam19, Cnn1, Col4a1, Ctgf, Ctps, Rflnb, Fstl3, Hspb1, Igfbp3, Pxdc1, Sema7a, Sh3pxd2a, Tagln, Tgfbi, Tns1, Tpm1)^5^.

##### Single-cell RNA-seq analysis

Single-cell RNA-seq fastq files were processed through the count utility from CellRanger (10X Genomics)^48^ using a custom reference generated from GENCODE M15 gene annotations and GRCm38/mm10. Multiplexed samples and antibody specific barcodes were parsed using a wrapper to the DemuxEM^49^ package. The resulting UMI counts were read into Seurat and TCR information was added to the metadata using scRepertoire^50^. Cells were filtered for singlets marked by DemuxEM. Cells were filtered for more than 500 genes expressed and less than 5% of reads from mitochondrial genes.

After each of the described filtering steps, data was processed as follows: Data was log-normalized using the NormalizeData function from Seurat^51^. The top 2,000 variable features were selected using FindVariableFeatures, and the data was scaled using ScaleData from Seurat. Principal components were determined using RunPCA, and nearest neighbors were calculated using FindNeighbors with 30 principal components (PCs). The data was clustered to a specified resolution using FindClusters with 30 PCs. UMAPs were generated with 30 PCs using RunUMAP. For the avb8 scRNA-seq study, cells were clustered to a resolution of 0.2 and three clusters containing stromal cells were removed. Cells were sorted into DC or T cells. The T cells were clustered at a resolution of 0.2, and one cluster of myeloid cells was removed. scGate^52^ was run using the built-in generic mouse models for CD8T and CD4T. Cells classified as CD4 T cells were kept, and TCR genes were removed to avoid biasing clustering. After reclustering to resolution 0.5, a cluster of gamma-delta T cells and a cluster of naïve T cells were removed. The CD4 T cells were reclustered to resolution 0.2, and four clusters of Tregs were kept.

The DCs were clustered to a resolution of 0.2, and seven clusters of lymphocytes, monocytes/macrophages and mast cells were removed. After reclustering to a resolution of 1, two clusters of NK and B cells were removed. Clustering was performed again to a resolution of 0.2 and two clusters of monocytes/macrophages were removed.

Pseudobulk differential expression was performed by averaging the expression of the cells per animal using the AverageExpression function from Seurat. Scoring for the GO term antigen processing and presentation pathway (GO:0019882) was done by averaging expression using AverageExpression from Seurat, then scoring the samples using ScoreSignatures_UCell from UCell^53^ .

Data in figure 1C is from^24^, reprocessed from fastq to be consistent with the pipeline versions used here. After the QC filtering described above, the cells were clustered to a resolution of 0.1 and two clusters containing T cells and fibroblasts in the epithelial sort were removed. scGate^52^ was run using the built-in generic mouse models for CD8T and CD4T. Only cells classified as CD4 or CD8 T cells were kept for the T cell compartment.

Tumor draining lymph node CD8 and CD4 T cells were processed in the same initial steps. After the QC filtering described above, TCR genes were removed to avoid biasing clustering. scGate^52^ was run using the built-in generic mouse models for CD8T and CD4T. Only cells classified as CD4 or CD8 T cells were kept. The CD8 T cells were clustered to a resolution of 1 and a cluster of Tregs was removed. CD8 T cells were clustered to a resolution of 0.8. CD4 T cells were clustered to a resolution of 0.2.

### Ex-vivo antigen presentation assay by lymph node DCs

Lymph nodes (inguinal, axillary, and brachial) from mice treated with anti-gp120 (control), anti-αvβ8 /anti-PDL1 combination, or anti-TGFβ/anti-PDL1 combination as described above were collected from mice 24 hour after treatment. Lymph nodes were digested into single cell suspensions using collagenase P, DNAse, and dispase as described above, and cells were stained for viability dye (1/1000, Ghost Dye Red 780, Cytek Biosciences, Cat no. 13-0865-T100) and Fc receptor-blocked (1/50, BD Biosciences, Cat no. 553142) in PBS for 20 minutes on ice. Cells were washed and resuspended in primary antibodies (1/150, anti-Ly6G (BioLegend, Cat no. 127614); 1/150, anti-CD19 (Cytek Biosciences, Cat no. 20-0193-U100); 1/150 anti-NK1.1 (BioLegend, Cat no. 108710); 1/150 anti-CD90.2 (BioLegend, Cat no. 105312); 1/100 anti-F480 (BioLegend, Cat no. 123116); 1/150 anti-Ly6C (BioLegend, Cat no. 128016); 1/200 anti-CD11b (BD Biosciences, Cat no. 563553); 1/150 anti-CD45.2 (BioLegend, Cat no. 109847); 1/150 anti-MHC class II (Invitrogen, Cat no. 11-5321-82); 1/100 anti-CD11c (BioLegend, Cat no. 117308) in PBS buffer with 0.5% bovine serum albumin + 2mM EDTA for 20 minutes on ice. Cells were washed, filtered through a 70μm filter, and bulk DCs (Live, CD45.2^+^, Dump^-^ (Ly6G, CD19, NK1.1, CD90.2, F480, Ly6C), MHC class II^+^, CD11c^+^, CD11b^+/-^) were isolated by FACS from the lymph nodes. 20,000 DCs were seeded into a 96 well U-bottom plate in 150μL of complete RPMI media (10% heat-inactivated fetal bovine serum, 2mM Glutamax, and 100U/mL Penicillin-Streptomycin) containing 10μg/mL of ovalbumin (InvivoGen, Cat no. vac-pova), or media alone for 2 hours at 37°C. To isolate ovalbumin-specific CD8^+^ T cells, spleens from female OT-1 mice (Jackson Laboratory, strain 003831) were collected, mashed through a 70um strainer, and single cell suspensions were stained with the Naïve CD8a^+^ T cell Isolation Kit (Miltenyi, Cat no. 130-096-543). Naïve CD8a^+^ T cells were isolated according to manufacturer’s instructions and stained with a 4μM solution of CellTrace Violet (Invitrogen, Cat no. C34557) in PBS. The labelled T cells were then resuspended in complete RPMI media containing 2mM Glutamax (Gibco, Cat no. 35050-061), 0.01mM HEPES, 1X MEM non-essential amino acids (Gibco, Cat no. 11140-050), 1mM sodium pyruvate (Gibco, Cat no. 11360-070), and 55μM 2-mercaptoethanol (Gibco, Cat no. 21985-023). After the 2h incubation period, the ovalbumin-containing media was removed from the DCs and 50,000 ovalbumin-specific CD8a^+^ T cells in 150μL supplemented media were co-cultured with the DCs for 3 days at 37°C. Co-cultures were collected, stained with viability dye and FcR-blocked as described above, and stained in primary antibodies (1/150 anti-CD11c (BioLegend, Cat no. 117310); 1/150 anti-CD8 (Invitrogen, Cat no. MA516759) for 20 minutes on ice. CellTrace Violet dilution was then assessed in ovalbumin-specific CD8a^+^ T cells by flow cytometry.

## List of Supplementary Materials

Fig S1 to S7 for multiple supplementary figures

## Acknowledgments

### Author contributions

Conceptualization: DS, AC Methodology: KW, YQ, DS, AC

Investigation: SM, NSF, JWL, ZZ, CW, HYM, XR, CC, YY, JZ, MT, MM, AC

Formal Analysis: KW, AC Data Curation: KW Resources: CC, HP, WL, JTK

Supervision: KML, RC, SJT, YW, IM, NRW, SM, YQ, DS, AC

Project Administration: AC

Writing – original draft: KW, DS, AC

Writing – review & editing: KW, HP, KML, SJT, IM, NRW, YQ, DS, AC

---

**SUPPLEMENTARY FIGURE S1.**
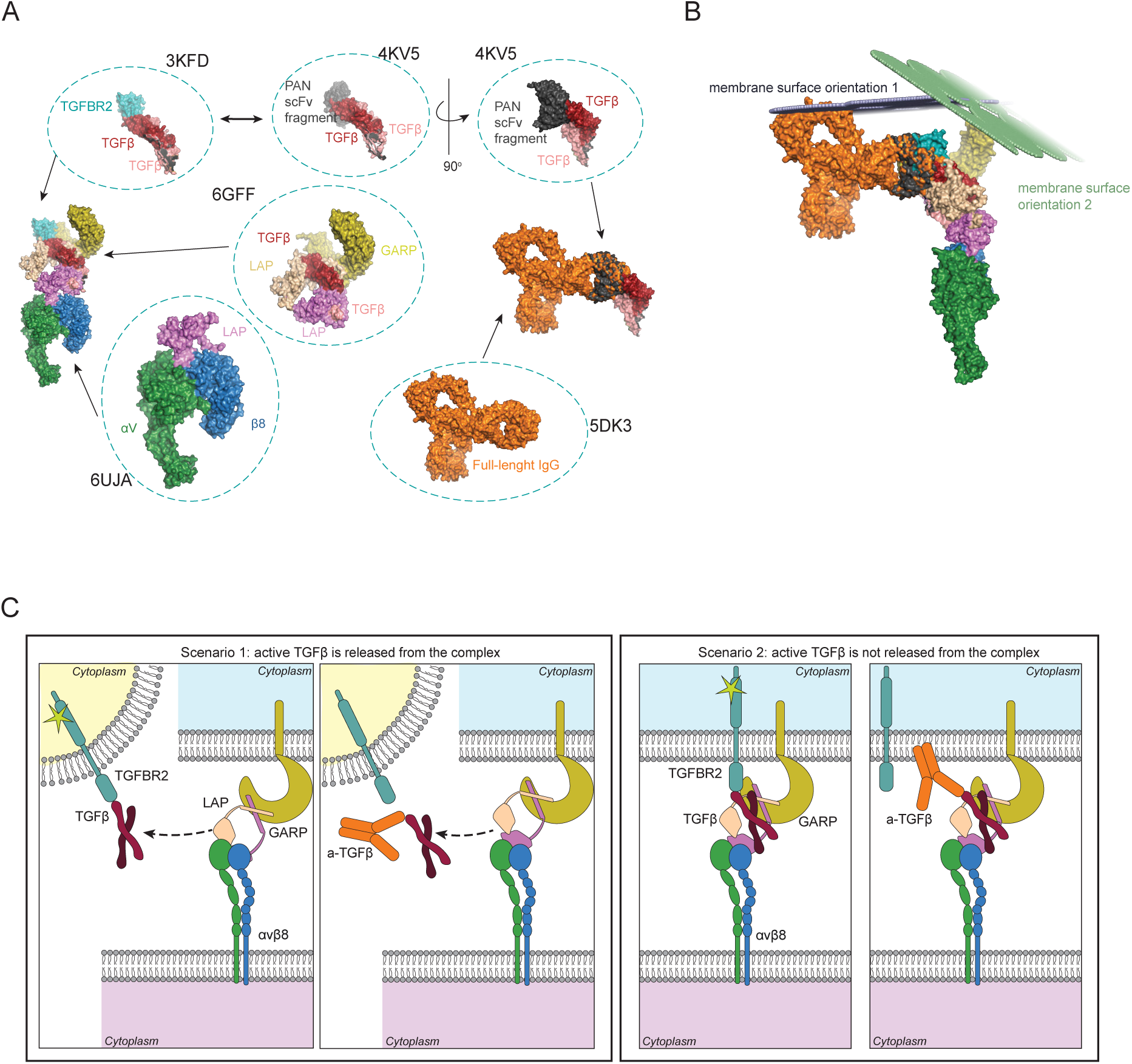
Structural model of the αvβ8-LTGFB1-GARP-TGFBR2 signaling assembly with anti-TGFβ antibody. (A) Composite structural model of the αvβ8-LTGFB1-GARP-TGFBR2 signaling assembly (leftmost panel), generated by overlaying experimentally determined structures of subcomplexes (shown in the dashed circles), annotated with their respective Protein Data Bank identifiers (PDB ID). The αv and β8 subunits are colored green and blue, respectively; the mature TGFβ and Latency-Associated Peptide (LAP) dimers are colored red & pink and magenta & wheat, respectively. GARP is colored yellow and TGFBR2 cyan. Structural overlay of Pembrolizumab and the PAN anti-TGFβ scFv antibody bound to TGFβ (rightmost panel). The experimental structure of the PAN anti-TGFβ scFv antibody (colored black) bound to TGFβ indicates its epitope. A breakdown of the experimental structures used for this overlay is shown in the dashed circles, each annotated with its respective PDB ID. (B) Cartoon representation of two relative membrane surface orientations (shown in blue and green spheres) that would be compatible with simultaneous insertion of TGFBR2 and GARP in the same cell surface membrane, showing potential steric hindrance for an anti-TGFB antibody to bind. (C) Cartoon depicting the different mechanisms of TGFβ activation and the hypothesized access that anti-TGFβ antibody would have depending on the scenario. In scenario 1 (left), active TGFβ is released from the LTGFB-GARP complex, and the anti-TGFβ antibody is able to prevent binding to TGFBR2. In scenario 2 (right), active TGFβ is not released from the complex, and therefore, the anti-TGFβ antibody has limited access to its epitope due to the proximity of the cell membrane.

**SUPPLEMENTARY FIGURE S2.**
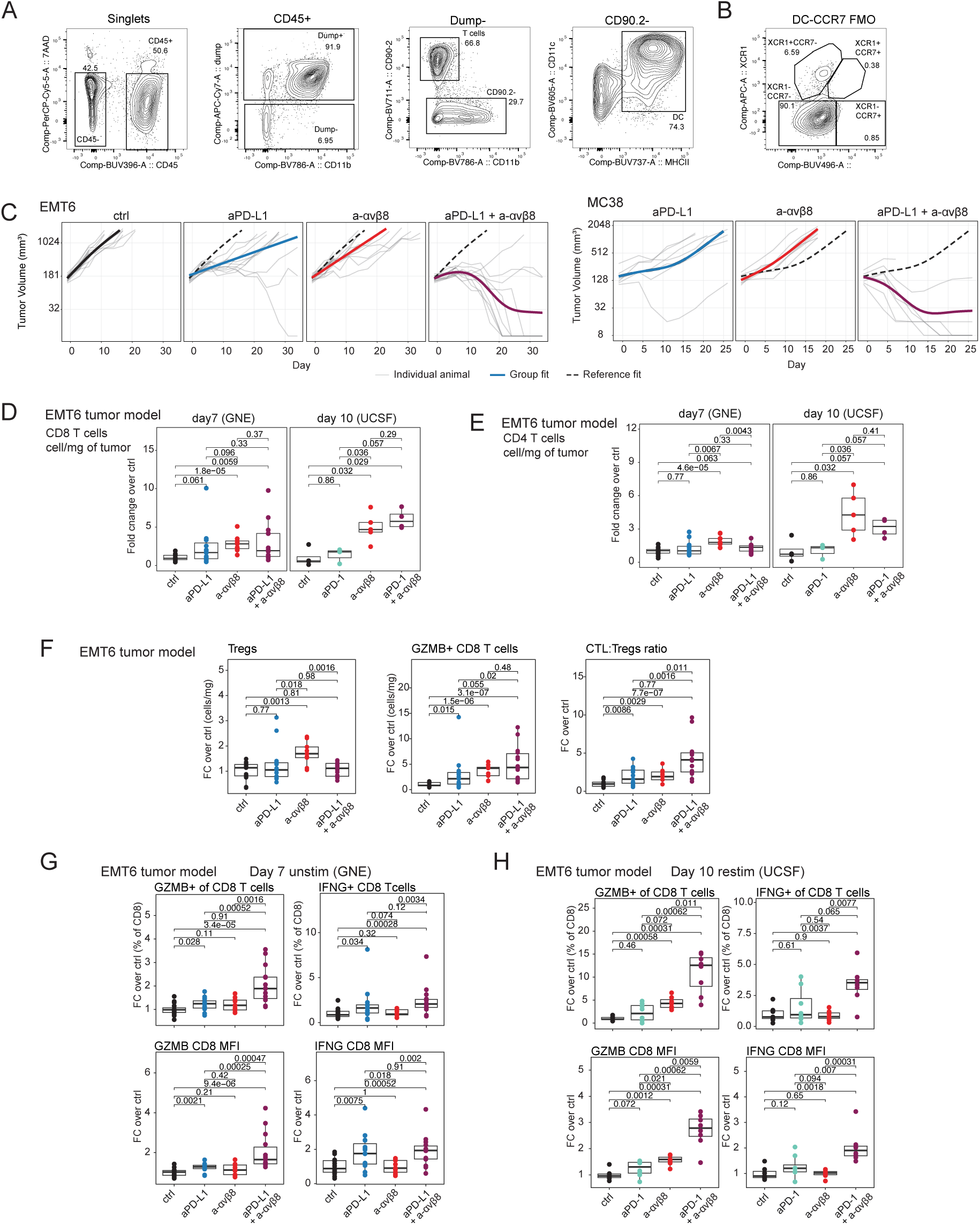
Related to Main Figure 2. Analysis of anti-tumor activity of anti-αvβ8/anti-PD-L1 combination treatment. (A) Gating strategy for the flow cytometry analysis shown in Figure 1D-F. Dump channel included: Ly6G, CD19, F4/80, and Ly6C. (B) FMO control for CCR7 staining in the DC gate. (C) EMT6 and MC38 tumor volume showing the individual animal curves relative to Figure 1G (n=10 animals per group). Flow cytometry quantification of (D) CD8 and (E) CD4 T cells infiltrating EMT6 tumors at day 7 (performed at Genentech) and day 10 (performed at UCSF) after initiation of treatment. Fold change over average of the control group (y axis; groups: x axis; data from: three independent day 7 experiments, three independent αvβ8 experiments, ctrl n = 15, aPD-L1 n = 15, a-αvβ8 n = 9, combo a-αvβ8 n = 14; one day 10 experiment, ctrl n = 4, aPD-1 n = 3, a-αvβ8 n = 5, combo a-αvβ8 n = 4). (F) Flow cytometry quantification of Tregs and GZMB+ CD8 T cells infiltrating EMT6 tumors at day 7 (performed at Genentech) after initiation of treatment. Fold change over average of the control group (y axis; groups: x axis; data from: three independent day 7 experiments, three independent αvβ8 experiments, ctrl n = 15, aPD-L1 n = 15, a-αvβ8 n = 9, combo a-αvβ8 n = 14). (G) Flow cytometry quantification of unstimulated GZMB+ and IFNg+ CD8 T cells infiltrating EMT6 tumors at day 7 (performed at Genentech) after initiation of treatment: percentages on the top, mean fluorescence intensity on the bottom. Fold change over average of the control group (y axis; groups: x axis; data from: three independent day 7 experiments, three independent αvβ8 experiments, ctrl n = 15, aPD-L1 n = 15, a-αvβ8 n = 9, combo a-αvβ8 n = 14). (H) Flow cytometry quantification of ex vivo stimulated GZMB+ and IFNg+ CD8 T cells infiltrating EMT6 tumors at day 10 (performed at UCSF) after initiation of treatment: percentages on the top, mean fluorescence intensity on the bottom. Fold change over average of the control group (y axis; groups: x axis; data from one day 10 experiment, ctrl n = 7, aPD-1 n = 8, a-αvβ8 n = 7, combo a-αvβ8 n = 8)

**SUPPLEMENTARY FIGURE S3.**
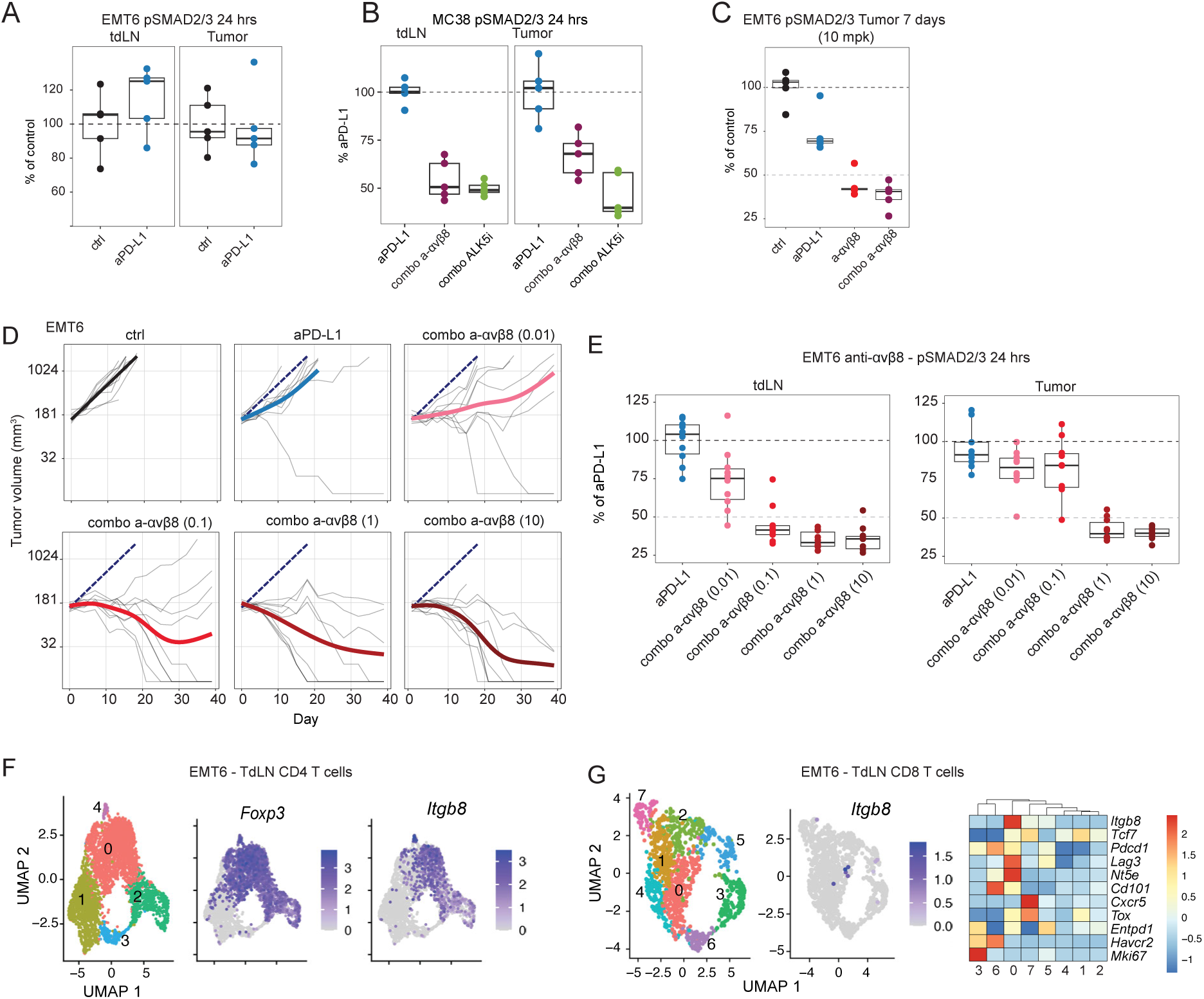
Related to Main Figure 3. Ex vivo assessment of TGFβ pathway inhibition. (A) *Ex vivo* assessment of SMAD2/3 phosphorylation (pSMAD2/3) by ELISA. Tumor draining lymph nodes (tdLN) and EMT6 tumors from mice treated with isotype control or anti-PD-L1 (24 hrs) (n=5 mice per group of treatment). (B) *Ex vivo* assessment of SMAD2/3 phosphorylation (pSMAD2/3) by ELISA. Tumor draining lymph nodes (tdLN) and MC38 tumors from mice treated with anti-PD-L1 alone or in combination with either an ALK5 small molecule inhibitor (1hr), or anti-αvβ8 antibody (24 hrs) (n=5 mice per group of treatment). (C) *Ex vivo* assessment of SMAD2/3 phosphorylation (pSMAD2/3) by ELISA. EMT6 tumors from mice treated with anti-PD-L1 alone or in combination with anti-αvβ8 antibody (7 days, one experiment, n=5 mice per group of treatment). (D) Tumor volume (y axis) of EMT6 tumors treated with a combination anti-PD-L1 with increasing doses of anti-αvβ8 (from 0.01 to 10 mg/kg) over time (x axis). Individual animal curves (grey lines) and group fit curves (thick solid lines) of the control group (black) and treatment groups (colored) are provided (n = 10 for all groups). (E) *Ex vivo* assessment of SMAD2/3 phosphorylation (pSMAD2/3) by ELISA. Tumor draining lymph nodes (tdLN) and EMT6 tumors from mice treated with anti-PD-L1 alone or in combination with increasing doses of anti-αvβ8 antibody (from 0.01 to 10 mg/kg, 24 hrs) (data from two independent studies, n=10 mice per group of treatment). (F) UMAP of 3,942 CD4 T cells (dots) sorted from untreated EMT6 tumor draining lymph nodes colored by cluster (n = 3 mice, left panel). *Foxp3* (middle panel) and *Itgb8* (right panel) expression levels [Log(CPM/100 + 1)] in UMAP space are shown. (G) UMAP of 1,153 CD8 T cells (dots) sorted from untreated EMT6 tumor draining lymph nodes colored by cluster (n = 3 mice, left panel). *Itgb8* (middle panel) expression levels [Log(CPM/100 + 1)] in UMAP space are shown. Heatmap of relative average expression of selected genes related to the T_PEX_ phenotype (*Tcf7+Pdcd1+ Lag3+Havcr2-*, cluster 0) in CD8 T cells populating different clusters (right panel).

**SUPPLEMENTARY FIGURE S4.**
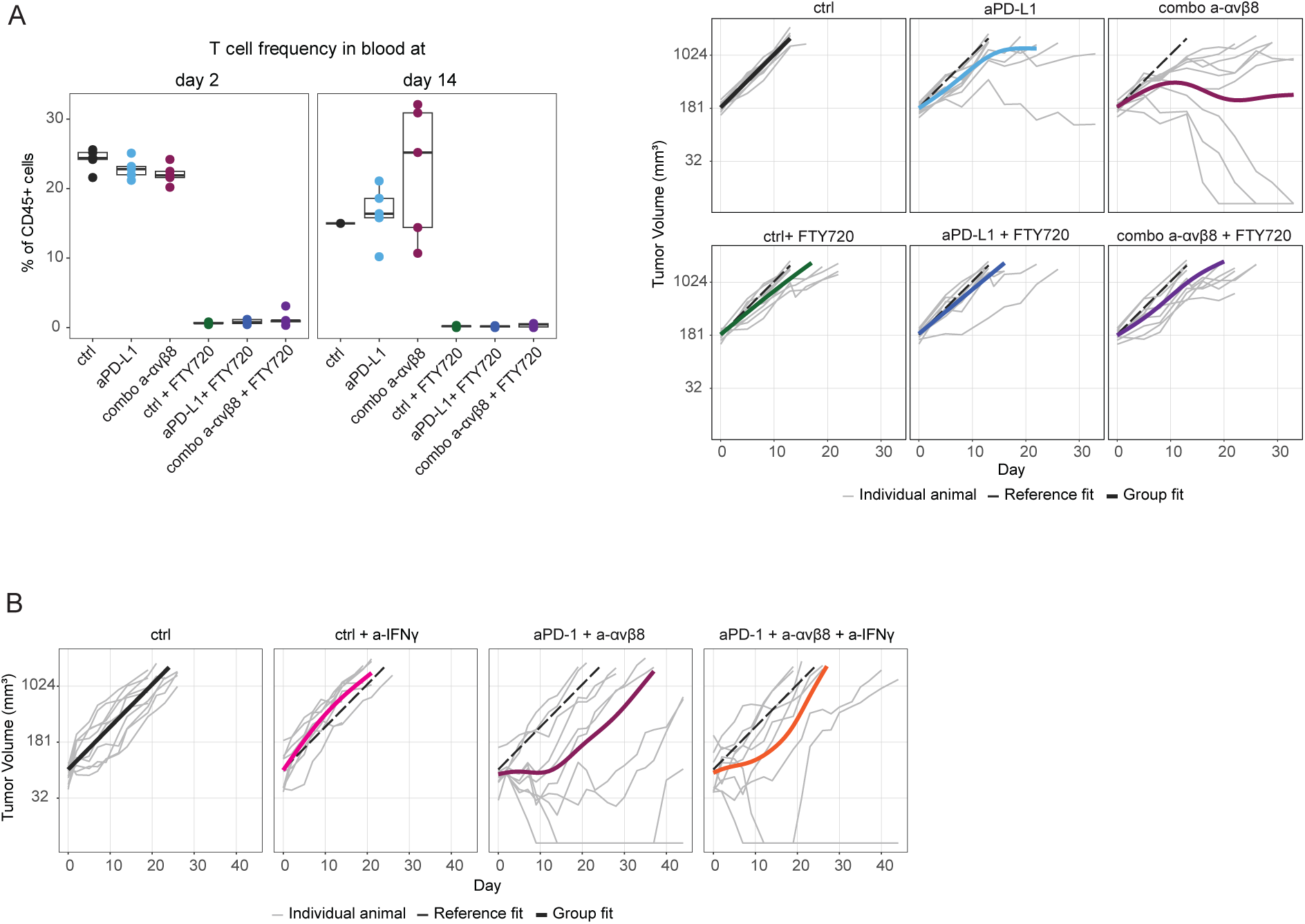
Related to figure 4. Additional experiments showing the anti-tumor activity of the anti-αvβ8/anti-PD-L1 combination in the presence of FTY720, and the anti-αvβ8/PD1 combination on top of IFNγ neutralization. (A) Left panel: Flow cytometry quantification of T cell frequency in blood from mice bearing EMT6 tumors treated with anti-αvβ8/anti-PD-L1 combination with or without FTY720 at day 2 (left) and day 14 (right) (n=5 per treatment group). Right panel: Tumor volume (y axis) of EMT6 tumors treated with anti-PD-L1/ αvβ8 combination with or without FTY720 over time (x axis). Individual animal curves (grey lines) and group fit curves (thick solid lines) of the control group (black) and treatment groups (colored) are provided (n = 10 for all groups). (B) Tumor volume (y axis) of EMT6 tumors treated with anti-PD-1/αvβ8 combination with or without anti-IFNγ over time (x axis). Individual animal curves (grey lines) and group fit curves (thick solid lines) of the control group (black) and treatment groups (colored) are provided (n = 9-10 for all groups)

**SUPPLEMENTARY FIGURE S5.**
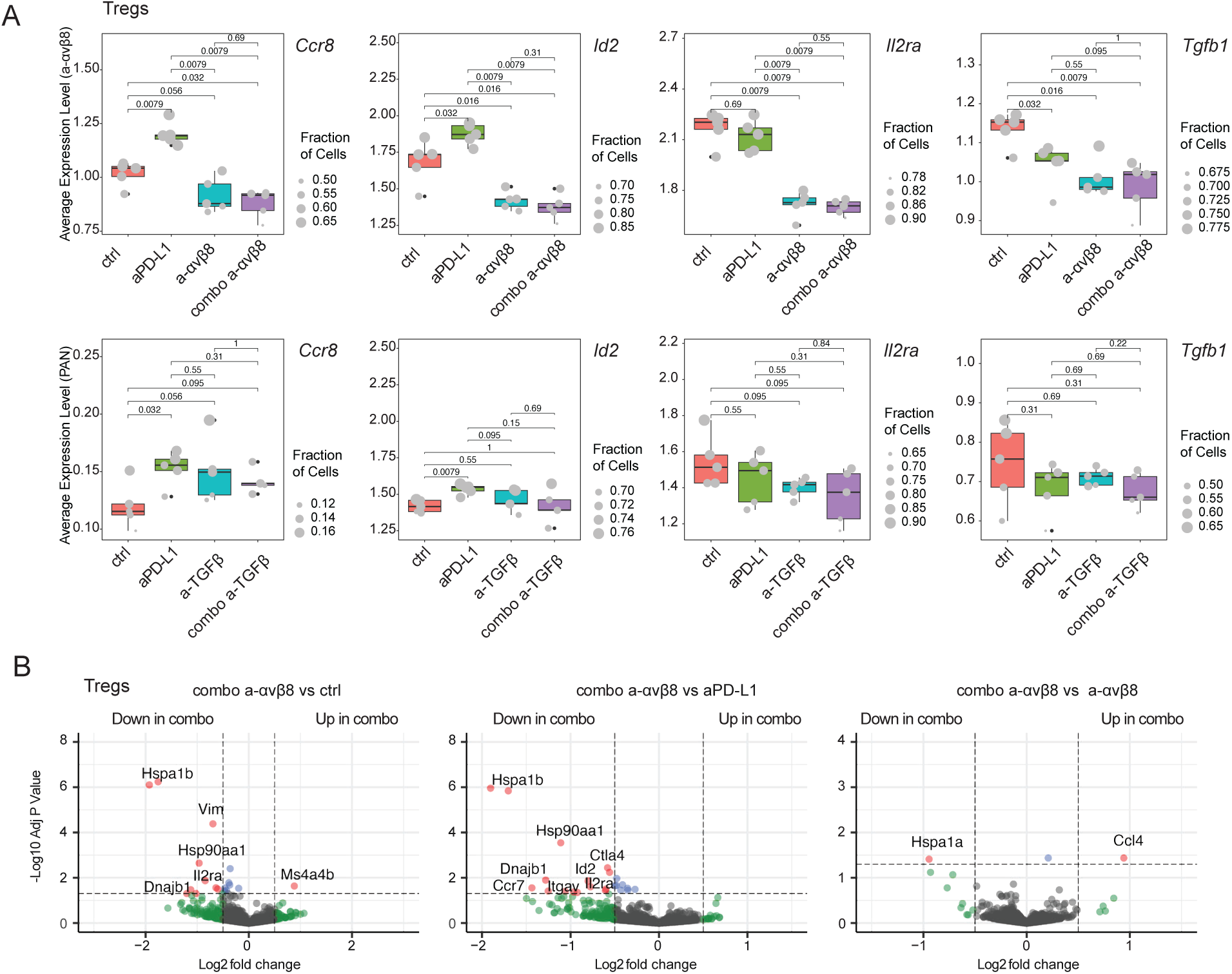
Transcriptomic analyses of the effect of anti-αvβ8 / PD-L1 anti-TGFβ/ PD-L1 combination on T_REG_s. (A) Expression of selected genes in the T_REG_ population in two independent experiments, EMT6 tumor-bearing mice treated with: anti-αvβ8 / PD-L1 antibodies (top panel) or anti-TGFβ/ PD-L1 combination (bottom panel). Individual animal values are shown, with dot size representing the fraction of T_REG_s expressing the specified gene (n=5 per treatment group). P values are from paired Wilcoxon rank-sum tests (two-sided) comparing treatment conditions. (B) Pseudo-bulk differential expression analysis comparing Tregs from the anti-PD-L1/ αvβ8 combination and either the control (left), the anti-PD-L1 (middle), or the anti-αvβ8 groups (n=5 per group). Dashed lines indicate significance thresholds: fold change, ± 0.5, and adjusted p-value< 0.05.

**SUPPLEMENTARY FIGURE S6.**
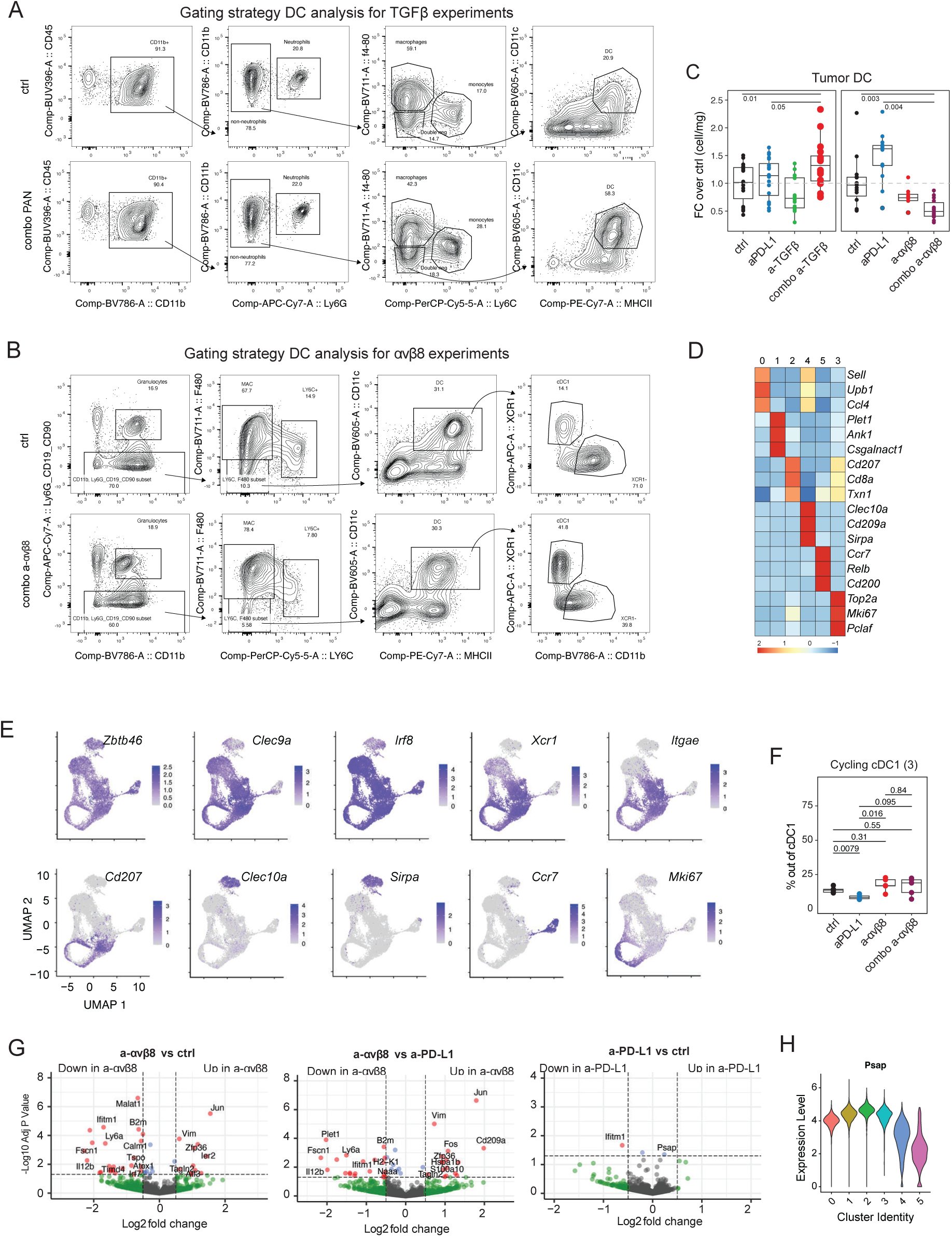
Related to main Figure 5. Analyses of tumor-infiltrating DCs. (A) Gating strategy for the flow cytometric analysis on EMT6 infiltrating DC in anti-PD-L1/ TGFβ experiments shown in Figure 4A. (B) Gating strategy for the flow cytometric analysis on EMT6 infiltrating DC in anti-PD-L1/ αvβ8 experiments shown in Figure 4A-B. (C) Flow cytometry quantification of EMT6 tumor infiltrating DCs (cells per mg of tissue) at seven days after initiation of treatment; anti-PD-L1/ PAN TGFβ combination (left), anti-PD-L1/ αvβ8 antibodies (right). Fold change over average of the control group (y axis; groups: x axis; data from: six independent PAN TGFβ experiments, ctrl n = 30, aPD-L1 n = 30, PAN n = 20, combo PAN n = 30; three independent αvβ8 experiments, ctrl n = 15, aPD-L1 n = 15, a-αvβ8 n = 9, combo a-αvβ8 n = 14). Stats: Dunn test with BH correction, adjusted P values of specific comparisons are shown. (D) Heatmap of relative average expression of three marker genes in each cluster from Figure 4C. (E) Expression levels [Log(CPM/100 + 1)] of selected genes in UMAP space from Figure 4C. (F) Quantification of the percentage of cells in cluster proliferating cluster 3 within the whole DC object (y axis; n = 5 per group of treatment from one experiment) in each animal (dots) by treatment group (x axis). Stats: Wilcox test with Holm correction; adjusted P-values are shown. (G) Pseudo-bulk differential expression analysis comparing EMT6 tumor-infiltrating DCs. Single agent comparisons are shown, anti-αvβ8 vs control (left), anti-αvβ8 vs anti-PD-L1 (middle), anti-PD-L1 vs control group (right) (n=5 per group). Dashed lines indicate significance thresholds: fold change, ± 0.5, and adjusted p-value< 0.05. (H) *Psap* expression across DC clusters, as identified in the UMAP shown in Figure 4C.

**SUPPLEMENTARY FIGURE S7.**
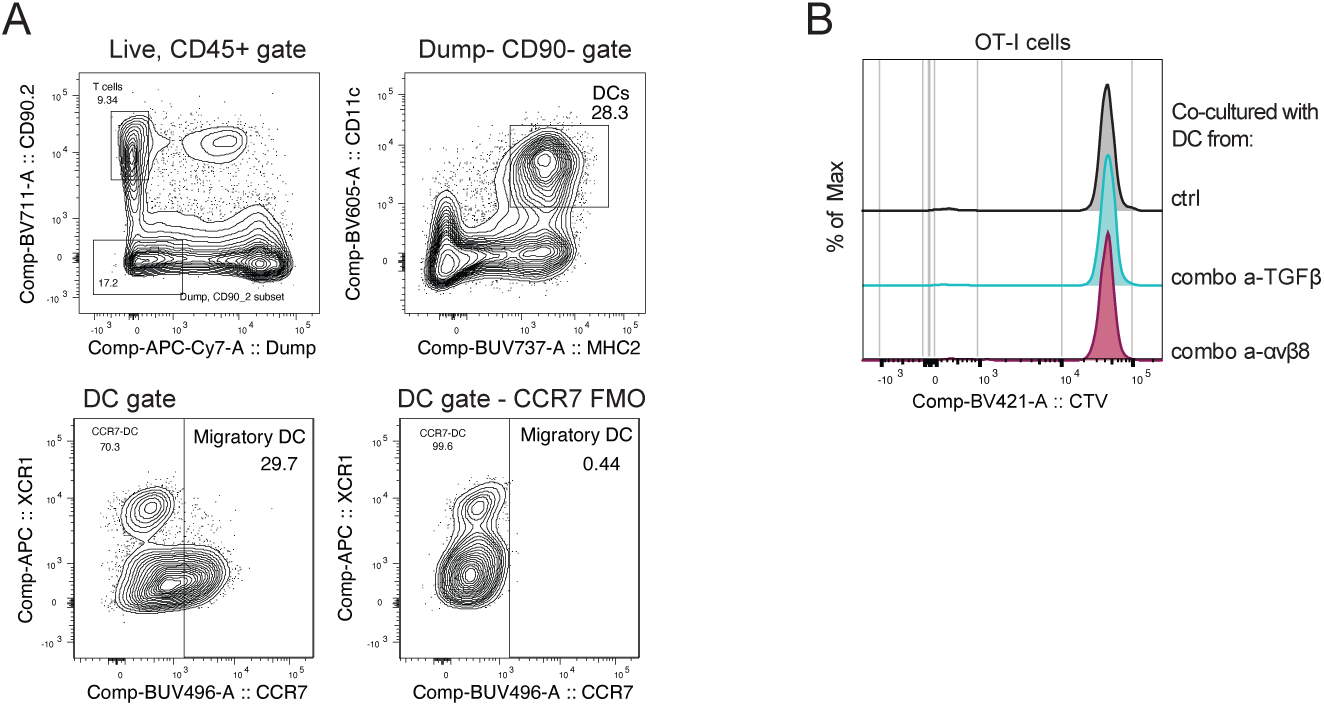
Related to main Figure 6. (A) Gating strategy and CCR7 FMO for the flow cytometric analysis of EMT6 infiltrating in figures 4F and G. Representative plots from an anti-PD-L1-treated mouse. (B) Flow cytometry analysis of OT-I cell proliferation based on the cell trace violet (CTV) fluorescence intensity. Representative plots of OT-I cells co-cultured for three days with DC from the lymph nodes of mice treated with control, anti-TGFβ/PD-L1, and anti-αvβ8/PD-L1 combination.

